# Sequence characterization of *T*, *Bip*, and *Phbw* demonstrates the role of MYB-bHLH-WD40 complexes and temperature in common bean seed color pattern formation

**DOI:** 10.1101/2023.09.23.559151

**Authors:** Travis Parker, Tayah Bolt, Troy Williams, R. Varma Penmetsa, Mwiinga Mulube, Antonia Palkovic, Celestina Nhagupana Jochua, Maria del Mar Rubio Wilhelmi, Sassoum Lo, Gail Bornhorst, Li Tian, Kelvin Kamfwa, Sam Hokin, Andrew Farmer, Christine Diepenbrock, Paul Gepts

## Abstract

Seed colors and color patterns are critical for the survival of wild plants and the consumer appeal of crops. In common bean, a major global staple, these patterns are also critical for determining market classes, yet the genetic and environmental control of many pigmentation patterns remains unresolved. In this study, we genetically mapped variation for three important seed pattern loci, *T*, *Bip*, and *p^hbw^*, which co-segregated completely with *PvTTG1*, *PvMYC1*, and *PvTT8*, respectively. Proteins encoded by these genes are predicted to work together in MYB-bHLH-WD40 (MBW) complexes, propagating flavonoid biosynthesis across the seed coat. Whole-genome sequencing of 37 diverse accessions identified putative mutations in each gene, including seven unique parallel mutations in *T* (*PvTTG1*) and a non-synonymous SNP in a conserved residue in *bip^ana^* (*PvMYC1*). A 612 bp intron deletion in *p^hbw^ (PvTT8)* eliminated motifs conserved since the origins of the Papilionoidea and corresponded to a 20-fold reduction in transcript abundance. Mutations in MBW candidate genes for *Z* (*PvTT2*) and *Sellatus* (WDR) were also identified. In multi-location field trials with seven varieties with partial seed coat patterning, pigmented seed coat area was highly unstable and correlated with temperature, with up to 11-fold differences in pigmented area between the warmest and the coolest environments. In controlled growth chamber conditions, an increase of 4 °C was sufficient to cause pigmentation on an additional 21% of the seed coat area. Our results shed light on the fundamental activation of flavonoid biosynthesis in common bean and will be instrumental for maximizing consumer appeal in this nutritious staple crop.

**Summary:** - Seed colors and patterns are critical for the survival of wild plants, and are important in differentiating crop market classes, but the genetic control of these in the staple crop common bean (*Phaseolus vulgaris*) is largely unknown.

- The genetic, transcriptional, and environmental basis of common bean seed color patterning was explored through QTL mapping, whole-genome sequencing, RT-qPCR, and automated pigmentation quantification of seed grown in multi-location field trials and growth chamber environments.

- MYB-bHLH-WD40 complex-forming genes *PvTTG1*, *PvMYC1*, and *PvTT8* co-segregated completely with the color patterning genes *T*, *Bip*, and *p^hbw^*. Mutations were identified in each gene, including seven unique parallel mutations in *T* (*PvTTG1*), a non-synonymous SNP in a conserved residue in *bip^ana^* (*PvMYC1*), and an intron deletion in *p^hbw^ (PvTT8)* eliminating highly conserved motifs and corresponding to 20-fold lower *PvTT8* transcript abundance. Mutations in MBW candidate genes *Z* (*PvTT2*) and *Sellatus* (WDR) were also identified. In multi-location field trials, pigmented seed coat area was highly unstable and corresponded to temperature. In growth chamber conditions, an increase of 4 °C caused pigmentation on an additional 21% of the seed coat area.

- Our results highlight the critical interaction between MYB-bHLH-WD40 complex components and temperature in establishing seed pattern diversity.

## Introduction

Proper specification of pigmentation in tissues such as flowers and seed is essential for the survival of wild plants, and is critical for market acceptance of crops. In the wild, pigmentation is vital for pollinator attraction, camouflage of seed to reduce herbivory, and promotion of seed dispersal. During domestication, a radical change in selection criteria for plant color patterns occurred and domesticates have undergone a nearly universal color diversification of harvested organs (Darwin, 1868, Hufford et al. 2019). Despite this, within most individual cultures and markets, a narrow range of preferred colors and color patterns exists and types must fall within this range to be culturally accepted and commercially successful.

Among grain legume crops, the common bean (*Phaseolus vulgaris* L.) is the most important species for direct human consumption (Broughton et al. 2003, Parker and Gepts 2021). Consumers and farmers exhibit strong preferences for dry beans with specific colors and patterns (as well as seed shape and size), which together account for the wide diversity of seed types found in common bean and related grain legumes. Diverse common beans with novel seed patterns have undergone a resurgence in developed countries. Consumers are willing to pay a premium for specialty dry beans (Walters et al. 2011, Brouwer et al. 2016, Swegarden et al. 2016), offering a market opportunity for their production.

Unlike seed color patterns related to the complex *C* locus on chromosome Pv08, partial coloring related to genes such as *T*, *Bip*, and *P* is often maintained after cooking. Seed coat color is primarily due to pigmented phenolic compounds. Phenolics (including flavonoids, phenolic acids, and proanthocyanidins) are one of the largest classes of specialized metabolites in plants, are biosynthesized in the shikimate/phenylpropanoid pathway, and in common bean among other grain legumes, are primarily localized in the seed coats (Gálvez Ranilla et al. 2007, Singh et al. 2017). Alongside these aesthetic considerations, phenolics have antioxidant properties and other beneficial health effects (Kondratyuk & Pezzuto 2004, Gálvez Ranilla et al. 2007, Yang et al. 2018, Rodríguez-Madrera et al. 2020, Bai et al. 2023). Despite these and other important roles *in planta*, the genetic and environmental control of phenolic pigment biosynthesis is poorly understood in *Phaseolus* beans.

Several genes influence the production of phenolic pigments in common bean. These can be classified into two major groups: a) factors controlling the spatial distribution of phenolic pigments, including their presence or absence in given areas; and b) genes modifying the composition (or visible color) of pigmentation uniformly throughout the seed coat. The first category consists of several genes, including *Pigment* (*P*; Shull 1907, Koinange et al. 1996, McClean et al. 2002, 2018). The *P* gene is dominant-acting and encodes a basic helix-loop-helix (bHLH) protein that is necessary for color expression in seeds and flowers of common bean (McClean et al. 2018). This gene is located on chromosome Pv07 and is orthologous to *Arabidopsis thaliana TRANSPARENT TESTA 8* (*AtTT8*, McClean et al. 2018). While full loss-of-function *p* alleles lead to complete loss of pigmentation in seeds and flowers, an allelic series of four partial loss-of-function alleles also exists at *P*, leading to spatially restricted pigmentation (Bassett 2007). This indicates that P is a spatial patterning gene, with full loss-of-function alleles affecting phenolics production across all seed and floral tissues. Full loss-of-function *p* mutations have arisen independently at least ten times (McClean et al. 2018).

Phenolic biosynthesis activation is also controlled by several other genes. The main factor for this is *Totally* colored (*T*), which interacts with other genes such as *Bipunctata* (*Bip*), *Zonal* (*Z*), and *Joker* (*J*) as the fundamental regulators of the presence or absence of pigment within any given cell of the seed coat. The fully recessive *t* allele eliminates floral pigmentation, and in the absence of secondary modifiers, may lead to a white patch of varying size opposite of the chalazal side of the hilum (the “*expansa*” pattern; Ernest et al. 2005, their Fig 2). All other partly colored patterns are achieved by *t* in combination with other mutations in *Bip, Z, J, Fib,* and/or partial loss-of-function alleles at *P* (Bassett 2007). Allelic series have been described for *Bip* (*Bip* > *bip^ana^* > *bip*) and *Z* (*Z* > *z^sel^*> *z*); with *bip^ana^* leading to the anasazi pattern and *z^sel^* leading to a yellow eye-type *sellatus* pattern. The chromosomes associated with each gene have been mapped through RAPD and STS markers (Brady et al. 1998, McClean et al. 2002, Reinprecht et al. 2013), but candidate genes for *T*, *Bip*, and *Z* are unknown, as are the molecular mechanisms of these genes and of the partial loss-of-function alleles at *P*.

In Arabidopsis, biosynthesis of phenolics, particularly flavonoids, anthocyanins, and proanthocyanidins, is under the control of MYB-bHLH-WDR (MBW) transcription factor complexes (Baudry et al. 2004, Xu et al. 2015, Zhang and Schrader 2017). Among the most important of these is the WDR-family gene *TTG1*, which combines with bHLH proteins such as *TT8* and *MYC1,* as well as numerous R2R3-MYB family transcription factors, including *TT2,* a critical flavonoid regulator (Baudry et al. 2004, Zhang and Schrader 2017). The complexes formed by these proteins further self-reinforce the expression of *TT8*, propagating the seed pigmentation signal from the chalaza through the rest of the seed coat (Baudry et al. 2006). These MBW complexes also activate expression of late biosynthetic genes (LBGs) related to phenolic production, including BANYULS (BAN, encoding anthocyanidin reductase; Baudry et al. 2004). Wu et al. (2019) and García-Fernández et al. (2021) characterized a cluster of *MYB113* homologs on Pv08 responsible for controlling the complex *C* pigmentation patterning locus of common bean. *MYB113* is an R2R3-MYB transcription factor and important MBW protein complex component; its homologs in cowpea control black seed color (Herniter et al. 2018). Similarly, the common bean seed color patterning gene *J* was recently linked to the R2R3-MYB transcription factor Phvul.010G130600 (Erfatpour et al. 2018, 2021; Erfatpour and Pauls 2020).

In this work, we seek to better understand the major regulatory factors influencing seed coat pigment biosynthesis in common bean. This entails the genetic mapping of several genes relevant to seed color in common bean, identification of interactions among these genes, whole-genome sequencing of 37 accessions with diverse seed color and patterns, gene expression analyses, and characterization of environmental effects on seed coat patterns in field research trials with growth chamber validation. Together, these results seek to answer fundamental questions about the control of seed patterning in common bean by interacting genetic and environmental factors.

## Materials and Methods

### Plant materials

The accessions ‘Black Nightfall’ (W6 51267) and ‘Orca’ (PI 632344, Hang et al. 2003) were used as parents in reciprocal crosses to develop a 217-member “BXO” recombinant inbred line (RIL) population (Parker et al. 2020). Black Nightfall has seed and floral patterns resembling the *half banner white* partial loss-of-function allele of *P* (*p^hbw^*, Bassett 1996, 2003). Orca is a progeny of Anasazi with a similar pattern, making it likely to bear *P t bip^ana^* (Bassett 2000). F_7_ plants were field-grown in Davis, California, in 2018, and F_8_ seeds were harvested and phenotypically classified into one of six phenotypic categories based on seed and flower color. A panel of 37 lines selected for seed color pattern diversity was used for whole-genome shotgun sequencing. The partly-colored varieties ‘Anasazi’ (PI 577705), ‘Orca’, ‘UC Southwest Gold’ (PI 693470, Parker et al. 2021a), ‘UC Southwest Red’ (PI 693472, Parker et al. 2021b), ‘UC Sunrise’ (PI 693474, Parker et al. 2021c), ‘UC Four Corners Red’ (PI 693469, T. Parker, unpublished), ‘UCD Jacob’s Cattle,’ and ‘UCD Holstein’ (both S. Temple, unpublished) were used to evaluate environmental impacts on seed color.

### QTL mapping and identification of candidate genes

For QTL mapping, DNA was extracted from individual F_8_ seeds of each line using a modified-CTAB protocol (Allen et al. 2006). DNA quality was checked by NanoDrop spectrophotometer and agarose gel electrophoresis and genotyped with the Illumina BARCBean6K_3 BeadChip (Song et al. 2015), yielding 1164 segregating single-nucleotide polymorphisms (SNPs). Linkage mapping was conducted using the ASMap R package (Taylor and Butler 2017, R Core Team 2022). QTL mapping was conducted on qualitative allelic states using maximum likelihood through the EM algorithm of the R/qtl package in R (Broman et al. 2003). The 95th percentile of 1000 random permutations of the phenotypic data was used as a significance threshold (LOD = 3.03). The phenotype and genotype data at the most significant SNPs were compared to identify SNPs flanking each seed coat patterning gene.

Descriptors and ontology information for all gene models between flanking SNPs of each QTL were extracted from Phytozome 13 (Goodstein et al. 2012; Schmutz et al. 2014; https://phytozome-next.jgi.doe.gov/info/Pvulgaris_v2_1). The amino acid sequences of Arabidopsis seed coat pigmentation regulators were obtained from TAIR and searched by BLASTp against the non-redundant proteomes of *P. vulgaris* and *A. thaliana*. Fast minimum evolution trees were developed based on the Grishin protein matrix on the NCBI website (<u>BLAST: Basic Local Alignment Search Tool</u> (nih.gov)) to compare patterns of sequence similarity and homology.

### Whole-genome sequencing and analysis

Whole-genome shotgun sequencing was conducted on the collection of 37 diverse accessions (Supporting Table S1) to better understand the sequence differences at candidate genes. DNA was extracted from seed embryonic axes of each variety using the Qiagen DNeasy Plant Minikit. Concentration and quality were checked by gel electrophoresis and Nanodrop, and the extracted DNA was sequenced by Illumina paired-end 150 bp sequencing at the UC Davis Genome Center. 270 Gb of sequence data were generated using a NovaSeq 6000 instrument, yielding an average of 12x sequencing depth per sample.

Sequence quality was checked with FASTQC (Andrews, 2010), and adapters and barcodes were trimmed using the Trimmomatic v0.39 (Bolger et al. 2014) ILLUMINACLIP function. Low-quality sequence was removed using the SLIDINGWINDOW:4:20 MINLEN:50 settings. Trimmed reads were aligned to the genomes of the Andean genotype G19833 v2.1 (Schmutz et al. 2014) and the Mesoamerican genotype UI111 v1.1 using the Burrow-Wheeler Aligner (BWA; Li and Durbin, 2009). Alignments were then imported and compared at regions of interest in the Integrative Genomics Viewer (IGV; Robinson et al. 2011). Predicted protein models were developed and visualized using the SWISS-MODEL Expasy web server [<u>SWISS-MODEL</u> (expasy.org)].

### Marker development and validation in recombinant lines

To evaluate the genotype of RILs with recombination, missing data, or heterozygosity between the markers flanking identified QTL at candidate mutations, DNA was extracted using a rapid NaOH-based protocol (Stonehouse et al. 2022, https://tinyurl.com/v24wzz4y) from Black Nightfall, Orca, and 45 BXO RILs. For three RILs with heterogeneous seed, DNA was extracted separately for each phenotypic class. These 47 lines were genotyped at all three loci to determine patterns of cosegregation between candidate mutations and phenotype. PCR and restriction digests were performed using the primers, cycling conditions, and restriction digest protocols in Supporting Table S2. PCR products were run on a 2.5% agarose gel for 25 minutes and visualized on a UV transilluminator to score indel size or CAPS cleavage at loci of interest.

### RT-qPCR and intron splicing

To quantify gene transcriptional patterns and evaluate patterns of intron retention or excision, RNA was sampled from developing seeds of greenhouse-grown plants at two time points: 10 days after anthesis (seeds approximately 5 mm long, 30 mg) and 20 days after anthesis (seeds approximately 10 mm long, 300 mg). Samples were flash-frozen in liquid nitrogen and maintained at -80 °C. RNA was extracted with the Qiagen RNeasy Plant Mini kit, with concentration and quality subsequently confirmed by NanoDrop and 1% bleach gel electrophoresis. Reverse transcription was conducted with the Invitrogen SuperScript IV VILO Master Mix and ezDNase kit. To evaluate the excision status of the second *TT8* intron, primers were designed to span from the second to the third exon (Supporting Table 2). PCR was performed on cDNA templates; the amplification product was run on a 2% agarose gel for 25 minutes and visualized on a UV transilluminator.

For RT-qPCR, transcript identifiers for each candidate gene were used to generate primers in NCBI primer BLAST with specificity checking to avoid amplification of non-target sequences. Intron-spanning primers were developed for all multi-exon genes, and controls without reverse transcriptase were run. RT-qPCR was conducted using the Qiagen Quantinova SYBR Green PCR Kit. The efficiency of qPCR primers was checked on pooled cDNA, and all selected primer pairs were shown to amplify with an efficiency of 0.96-1.08. The difference in C_T_ values between candidate and reference genes was used to generate ΔC_T_ data, which was then logarithmically converted to 2^-ΔC_T_ values. Differences in expression between phenotypic groups were then compared by anova and LSD.test() in the agricolae R package (de Mendiburu 2019).

### Environmental variables

To test for environmental factors affecting seed color, seven partly-colored varieties were field-grown at four locations across a coastal vs. inland climatic gradient in California, USA. These field sites were located in Pescadero (July average high/low: 21/11 °C; 37.223 °N, 122.359 °W; grown 2019), San Juan Bautista (27/12 °C; 36.847 °N, 121.535 °W; grown 2018), Valley Center (30/16 °C; 33.277°N, 117.026°W; grown 2018), and Davis (34/14 °C; 38.541 °N, 121.767 °W; grown 2019). For all experiments, a randomized complete block design was used with three replicated plots of 120 plants each. Seeds of each plot were bulk-harvested at dry maturity and approximately 100 seeds per lot were scanned to quantify the percent of pigmented seed coat area (PPSC). Scans were conducted using a Brother (Nagoya, Japan) DCP-7065DN scanner. Custom macros were developed in ImageJ to quantify the PPSC and total seed coat area. The quotient of these values was used to quantify the PPSC. The source code for the ImageJ macro is available at https://github.com/TravisParker91/Seed-color. Differences in PPSC were compared by two-way ANOVA with genotype and location as predictors and with an interaction term, and the LSD.test() in the agricolae R package was used with a Bonferroni correction for comparison of group means.

To test the isolated role of temperature on seed pigmentation, four plants of each of three partly colored varieties with compact growth habit were grown in two identical growth chambers of different temperatures, at 24/20 °C and 28/24 °C. Seeds were harvested from dry pods, and seeds were scanned to quantify pigmentation and statistically analyzed according to the methods used for field-grown seeds.

## Results

### Genetic component of seed pigment distribution

The BXO RIL population displayed transgressive segregation for seed and flower color, with seeds ranging from entirely black to almost entirely white, and flowers ranging from solid purple to white. In total, six unique seed/floral color classes were identified in the population. The predicted allelic combinations of Black Nightfall (*p^hbw^ T Bip*) and Orca (*P t bip^ana^*) were assembled into a potential genetic segregation model (Fig. 1) based on known specific effects of the three genes (Bassett 2007) and predictions of previously unstudied allele combinations (*p^hbw^ t bip^ana^*; *p^hbw^ T bip^ana^*). These combinations fit expected frequencies for the proportion of RILs in each category, supporting the model (χ^2^ test: *p*=0.14).

**Fig. 1.**
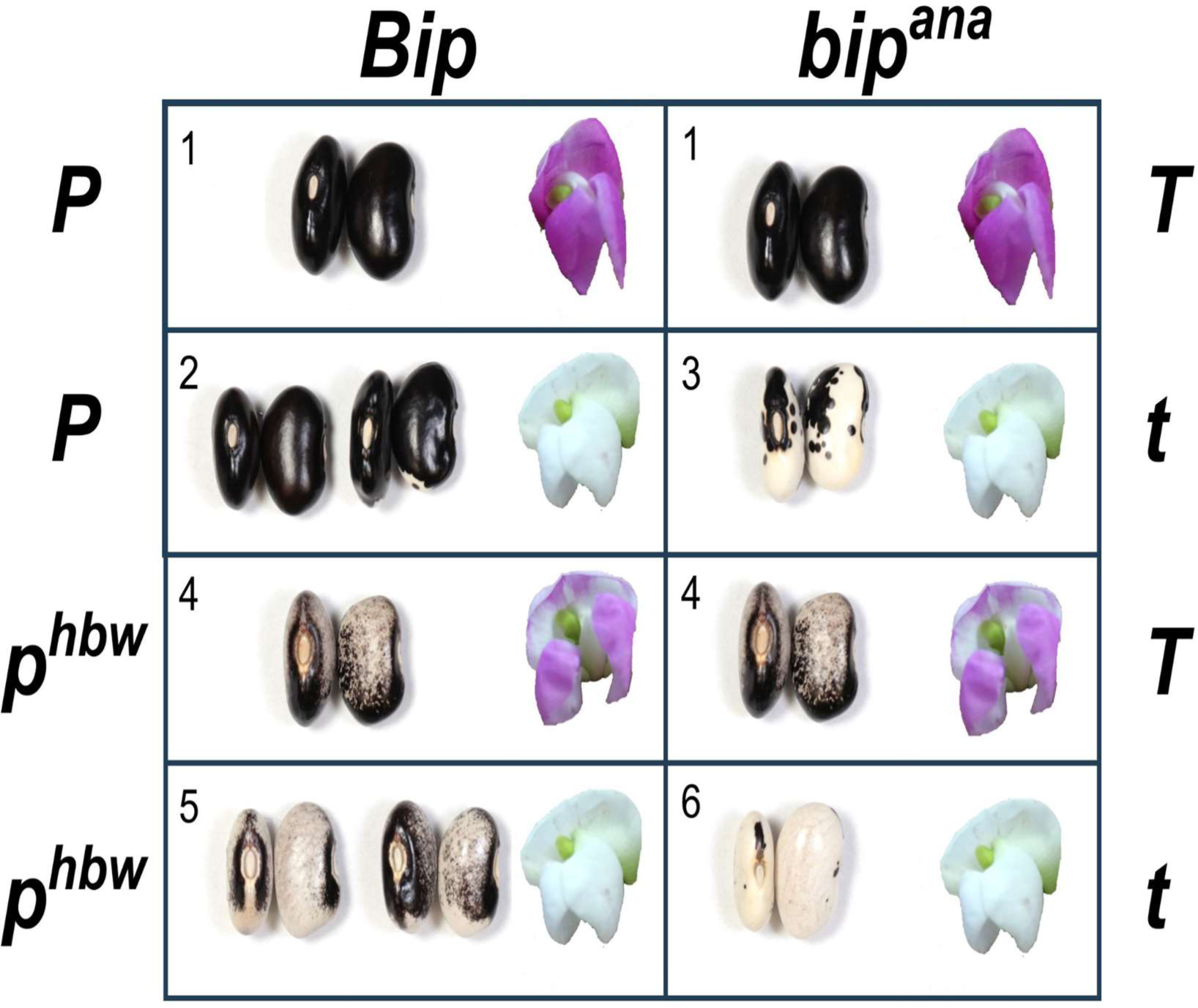
Genetic model of seed and floral patterning in the Black Nightfall/Orca (BXO) recombinant inbred line (RIL) population of common bean (P. vulgaris) based on the segregation of three loci with two alleles each: 1) P, phbw; 2) Bip, bipana; and 3) T, t. Six main phenotypic categories were observed in the population. The parental phenotypes Black Nightfall and Orca correspond to segregation classes 4 and 3, respectively, and the population shows transgressive segregation beyond this range of seed color patterning.

Significant QTLs for *p^hbw^, T,* and *Bip* were identified in the BXO population (Fig. 2). *T* mapped to chromosome Pv09 between 8,427,110 and 9,220,699 bp (*P. vulgaris* G19833 genome v2.1, Schmutz et al. 2014) (Table 1). This region included 63 predicted gene models, including the candidate gene Phvul.009G044700, a WDR protein whose closest Arabidopsis relative is *TTG1* (Supporting Fig. S1), a major seed coat color regulator. Phvul.009G044700 was therefore designated *PvTTG1*. Nine RILs showed recombination between the flanking markers, and the SNP closest to *PvTTG1* (at 283 kb in physical distance) was also the SNP most closely associated with floral and seed color patterns influenced by *T* (LOD = 53.8).

**Fig. 2.**
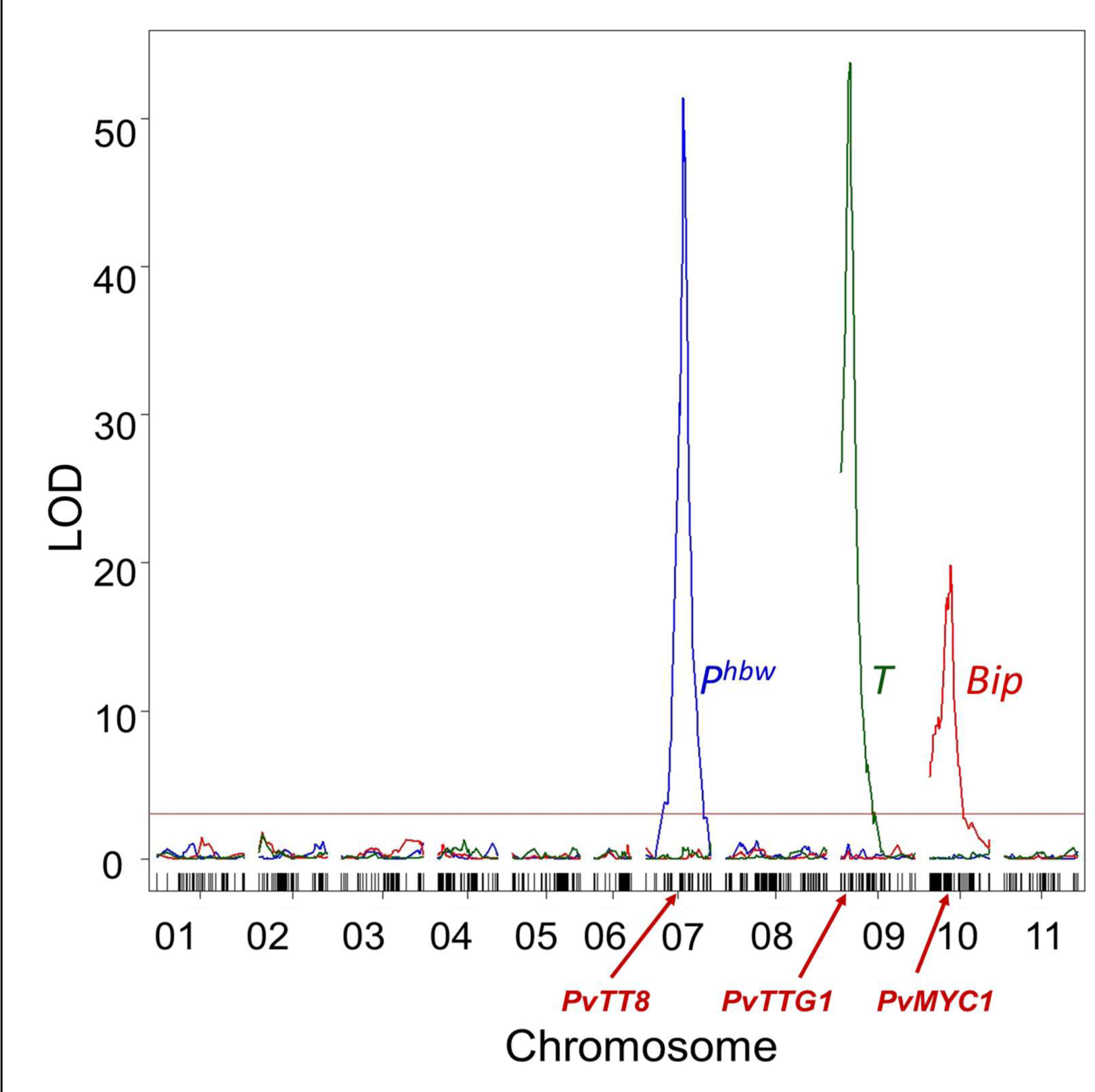
QTL mapping of the three major pigmentation-related genes (P, T, and Bip) in the Black Nightfall x Orca recombinant inbred population based on the percentage of pigmented seed coat area (PPSC). In each case, the most significant SNP is also the marker closest in physical space to the putative gene. The horizontal red bar is the significance threshold (LOD=3.03) obtained by permutation tests. The red arrows point to the location of the candidate genes for the three genes.

**Table 1.** Proposed unique mutant alleles with sequences characterized for the first time in this study.

| Allele | Arabidopsis homolog | Proposed gene model<sup>1</sup> | Variant | Example germplasm | Associated QTL (significant interval, bp) |
| --- | --- | --- | --- | --- | --- |
| *t<sup>Orca</sup>* | *TTG1* | Phvul.009G044700 | 39 bp deletion (del23_61) | Orca | Pv09: 8,427,110-9,220,699 |
| *t<sup>SYE</sup>* | *TTG1* | Phvul.009G044700 | 5 bp deletion (del422_426) | Steuben Yellow Eye |  |
| *t<sup>Rockwell</sup>* | *TTG1* | Phvul.009G044700 | 1 bp deletion (del749) | Rockwell |  |
| *t<sup>VA</sup>* | *TTG1* | Phvul.009G044700 | 1 bp deletion (del131) | Vermont Appaloosa |  |
| *t<sup>Snowcap</sup>* | *TTG1* | Phvul.009G044700 | SNP (730G>A) | Snowcap |  |
| *t<sup>DG</sup>* | *TTG1* | Phvul.009G044700 | Whole gene deletion | Dapple Grey |  |
| *t<sup>cf</sup>* | *TTG1* | Phvul.009G044700 | UTR 14 Mb inversion | Jacob's Cattle Gold |  |
| *bip<sup>ana</sup>* | *MYC1* | Phvul.010G098500 | SNP (650G>T) | Orca | Pv10: 36,099,863-36,771,237 |
| *p<sup>hbw</sup>* | *TT8* | Phvul.007G171333 | 612 bp deletion | Black Nightfall | Pv07: 28,312,574-29,971,196 |
| *p<sup>Hystyle</sup>* | *TT8* | Phvul.007G171333 | Deletion of first two exons | Hystyle |  |
| *z* | *TT2* | Phvul.003G132100 | 730G>A | Jacob's Cattle Gold | 1.4 cM from 31,468,000; McClean et al. 2002 |
| *sel* | WDR | Phvul.001G192200 | Promoter deletion at (-149 to -624 before 5'UTR) | Steuben Yellow Eye | 1.03 cM from 44,856,138; Bassett 1997 |

*Bip* mapped to chromosome Pv10 between 35,632,623 and 36,893,960 bp, a region with 76 predicted genes. Among these was Phvul.010G098500, predicted to encode a bHLH protein. Its protein product clusters as a sister clade to *MYC1* of Arabidopsis among all *A. thaliana* and *P. vulgaris* proteins (Supporting Fig. S2). Phvul.010G098500 was therefore named *PvMYC1*.

In Arabidopsis, *MYC1* is known to form complexes with *TTG1* to specify seed coat color. The SNP most strongly associated with *Bip* was found at 36,893,960 bp and was also the closest SNP in physical distance to *PvMYC1* (LOD=19.9). A MYB transcription factor Phvul.010G096400 was also found between the flanking markers. LOD scores for the QTL containing *Bip* were lower than those for the QTL containing *T* or *P*. This is due to the hypostatic nature of the gene relative to *T* (Fig. 1) as *Bip* could only be scored when *T* was homozygous recessive (Figs. 1, 2). Five RILs carrying a homozygous recessive *t* locus showed recombination between the flanking SNPs, and these were later screened for fine mapping at the candidate mutation.

The *p^hbw^* allele mapped to chromosome Pv07 between 28,312,574 (LOD=51.4) and 29,971,196, with 116 gene models and 16 gene models in the intervening space. This location is consistent with the demonstrated allelism of *p^hbw^* (formerly *stp^hbw^*) and *P* (Bassett 2003), which has previously been mapped to Phvul.007G171333 (*PvTT8,* McClean et al. 2018), a gene model in this interval. Each of the identified candidate gene models are MBW complex members (Fig. 3).

**Fig. 3.**
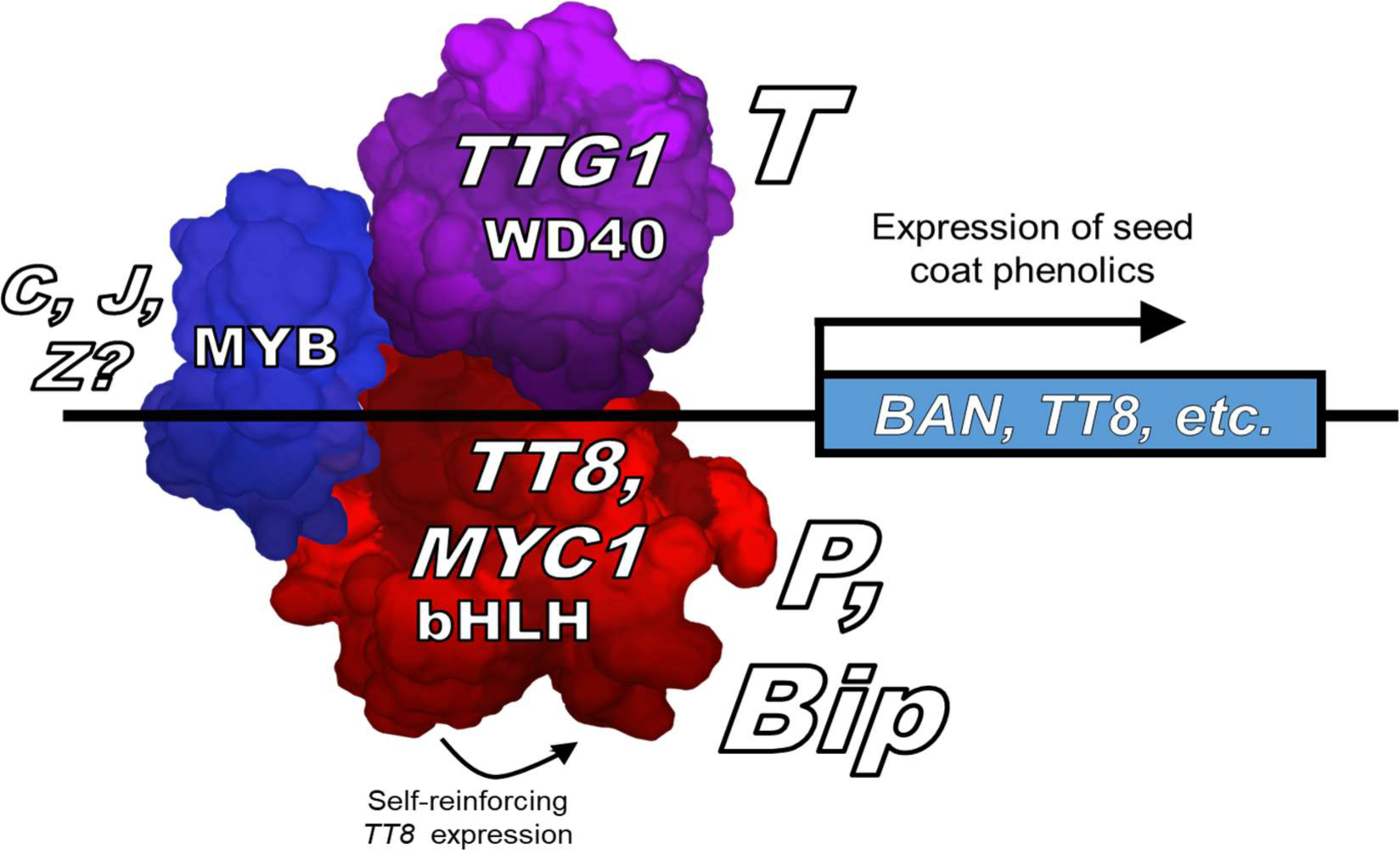
Proposed molecular genetic model for MYB-bHLH-WD40 (MBW) complexes affecting the expression of flower and seed pigmentation genes in Phaseolus vulgaris (see also Fig. 1 as example). Mapped genes (T, P, Bip, and potentially Z, J, and C) are predicted to encode proteins that form MYB-bHLHWD40 (MBW) complexes to activate flavonoid biosynthesis. In Arabidopsis, these require the inclusion of the WD40 protein TTG1, bHLH proteins such as TT8 or MYC1, and any of several R2R3-MYB transcription factors. PvTTG1, PvTT8, and PvMYC1, are found between the QTL flanking markers for T, P, and Bip, respectively.

### Sequencing

Sequence analysis in the 37 varieties (Fig. 4) at *PvTTG1* identified seven unique mutations that could explain loss of function in *T.* All partly-white seed types had one of these seven mutations, whereas none of the mutations were found in any fully-colored lines. The mutations included 1) a 39-bp deletion found in Orca and all five other Middle American partly-colored lines (*t^Orca^*), 2) a 5-bp deletion found in Steuben Yellow Eye (*t^SYE^*) and two other lines, 3) a single-base-pair deletion found in Rockwell (*t^Rockwell^*) and two other types, 4) a different single-base-pair deletion found only in Vermont Appaloosa (*t^VA^*), 5) a 1-bp, non-synonymous substitution in Snowcap (*t^Snowcap^*), 6) a 14-Mb inversion cutting into the 3’ untranslated region (UTR) of UCD Holstein and five other kidney-shaped lines with colored flowers (*t^cf^*), and 6) a deletion of the entire gene in Dapple Grey (*t^DG^*).

**Fig. 4.**
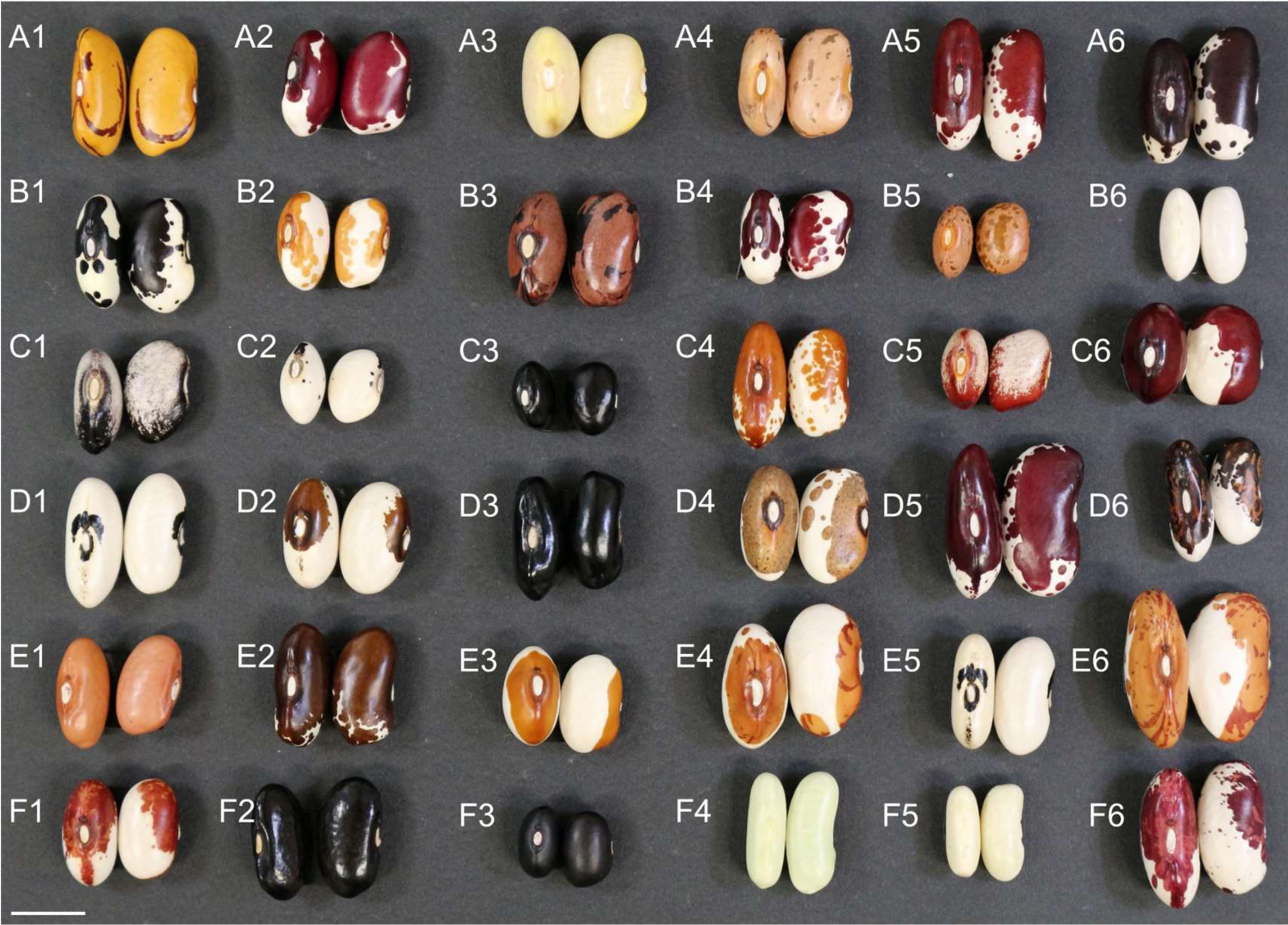
Seed coat diversity among 36 accessions used for whole-genome sequencing. Varieties were sampled from both domesticated gene pools and across most major economic classes, with a particular emphasis on types with partially white seed coats. For each accession, two seeds are shown, displaying the hilum side and the other showing the lateral side, with the caruncula or chalaza oriented above the hilum. Partly colored seed coats are controlled by interactions between T and other genes such as Bip, Z, J, and certain alleles at P. A 37th accession (not pictured) was an isogenic line of entry F5 with the same seed type. A1: UC Tiger’s Eye; A2: UC Four Corners Red; A3: UC Canario 707; A4: Bill Z; A5: UCD Jacob’s Cattle; A6: UCD Holstein; B1: Orca; B2: UC Southwest Gold; B3: UC Rio Zape; B4: UC Southwest Red; B5: MCM5001; B6: Vanilla; C1: Black Nightfall; C2: BXO075; C3: OXB178; C4: Jacob’s Gold; C5: Rio Colorado; C6: Red Calypso; D1: Improved Kidney Wax; D2: Davis; D3: Prolific Black Wax; D4: Dapple Grey; D5: Prefontaine; D6: Vermont Appaloosa; E1: UCD Pink 9634; E2: Pawnee Shell; E3: Steuben Yellow Eye; E4: Hidatsa Shield; E5: Burpee’s Kidney Wax; E6: Snowcap; F1: Rockwell; F2: Best of All Stringless Wax; F3: UI-911; F4: Hystyle (N3); F5: Prevail (I23); F6: Great Lakes Special. A 37th accession (not pictured) was an isogenic line of Prevail with the same seed type. Scale bar represents 1 cm.

The in-frame 39-bp *t^Orca^* deletion found in Middle American lines leads to elimination of residues 9-21 of the predicted protein product. These are found in one of the protein’s WD40 domains, which spans amino acid positions 9-54. Seven of the 13 amino acids deleted are conserved in alignments with *TTG1* in Arabidopsis. The three frameshift mutations (*t^VA^*, *t^SYE^*, *t^Rockwell^*) all led to premature stop codons. Whereas the wild-type amino acid chain has 336 amino acids, 254 are produced in *t^Rockwell^*, 177 in *t^SYE^*, and 94 in Vermont Appaloosa (Fig. 5). The 730G>A (D244N) substitution in *t^Snowcap^* affected a highly conserved residue, as Asp was the only residue found at this position among the 100 most similar proteins in the NCBI reference protein database, which included species of 21 plant families, indicating strong conservation and deleterious effects of substitution. A SIFT (https://sift.bii.a-star.edu.sg/www/SIFT_seq_submit2.html) analysis predicted that the D244N substitution would not be tolerated, and that Asp was the only amino acid residue not predicted to negatively impact gene function at the site. Types with the *T^cf^* allele showed a sequence break in the 3’ UTR, with mate sequences on both sides of the junction pairing with a similar break 14 Mb downstream on the chromosome, consistent with a major 14 Mb inversion. In Dapple Grey, *PvTTG1* was deleted in its entirety, with the deletion spanning from 15,507 bp upstream to 18 bp downstream of the gene.

**Fig. 5.**
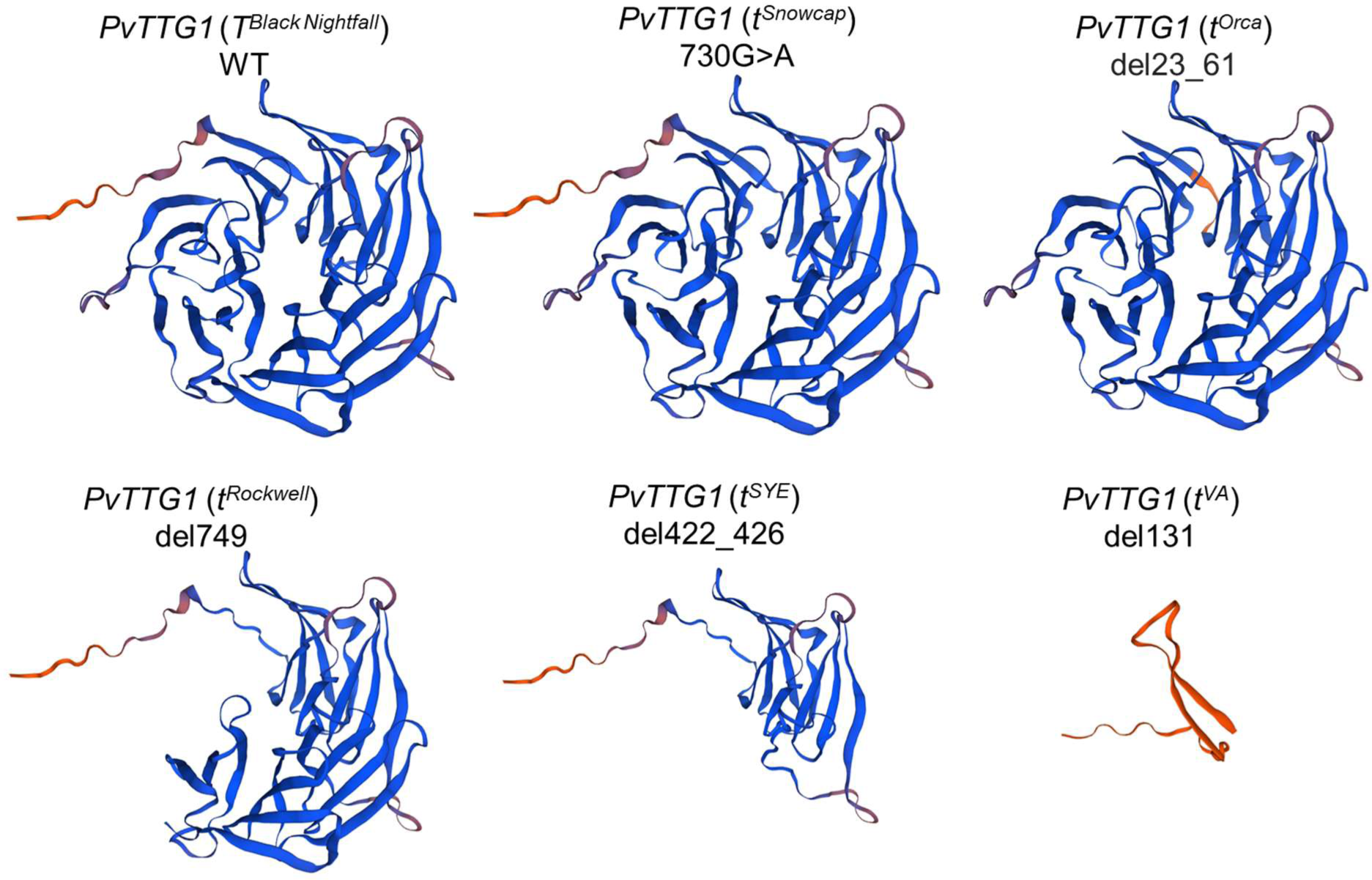
Structural variation in PvTTG1 (T) wild-type and mutant forms. A screen of 21 varieties with partly-colored seeds identified seven distinct parallel mutations in PvTTG1 (Phvul.009G044700). These include a non-synonymous substitution tSnowcap, two separate deletions of a single base pair (tRockwell and tVermont Appaloosa = tVA), a 5 bp deletion (tSteuben Yellow Eye = tSYE), a 39 bp deletion (tOrca), a 14 Mb inversion spanning into the UTR (tcf, not shown), and a deletion of the entire gene (tDapple Grey = tDG, not shown). Models were developed and visualized with the SWISS-MODEL Expasy web server (swissmodel.expasy.org).

Sequence evaluation of Phvul.010G098500 (*PvMYC1*) determined that Black Nightfall carried a race Mesoamerica haplotype at the locus, which differed from the race Durango haplotype carried by all five types with the Orca pattern. To identify polymorphisms related to *bip^ana^*, these types were therefore compared against the sequence of the race Durango breeding line UCD Pink 9634, known to also have wild-type *Bip* (Parker et al. 2021a,b,c). The *PvMYC1* coding sequence, UTRs, and 500 bp upstream of the transcription start site (TSS) only differed between wild-type UCD Pink 9634 and the mutant lines at two substituted base pairs. One of these, 1640C>T, led to an A547V substitution in a poorly conserved site. Seven different amino acids and a deletion occurred at this position across the 100 most similar protein models in the NCBI reference protein database. In contrast, the second SNP, 650G>T, leads to a non-synonymous C217F substitution at a highly conserved residue. Of the 100 most similar protein models in the NCBI reference protein database, representing nine plant families, 98 had cysteine residues at this site. The two exceptions were alternately spliced isoforms of a single soybean gene (Glyma.03G009500) which maintains the Cys in its primary transcript. No proteins in the database showed a non-synonymous substitution or deletion of the nucleotides encoding the Cys at position 217, indicating that the 650G>T substitution is highly likely to impact gene function. In contrast to *PvMYC1*, Phvul.010G096400 contained no non-synonymous SNPs or variation within 500 bp upstream of the TSS that distinguished Black Nightfall and Orca.

The three varieties displaying *p^hbw^* phenotypes (Black Nightfall, Rio Colorado, and the triple-mutant RIL BXO075) shared a 612 bp deletion from positions 19-630 bp in the second intron of Phvul.007G171333. This deletion was not identified in any other accessions in the panel. To identify and assess conserved regulatory motifs in the area, sequence data from the closest homologs of the gene across all legume species with above-ground seeds in the Phytozome database were extracted. BLAST sequence alignments identified motifs in the intron region that were conserved across all species analyzed (Supporting Fig. S3), except for the non-papilionoid legume *Cercis canadensis* (Koenen et al. 2021). The conservation of these intron motifs across all papilionoids, including those of the cool-season inverted repeat-lacking (IRLC) clade and the basal species *Lupinus albus*, contrasted with what was seen in other introns, which were weakly conserved. Intriguingly, homologous introns in the partly-colored reference lima bean accession G27455 and partly-colored reference cowpea accession IT97K-499-35 also display deletions in overlapping areas, where other members of each species show the full intron with conserved motifs (Supporting Fig. S3). The three sequenced varieties of snap (green) bean also shared a unique and previously undescribed full loss-of-function mutation at *P*, consisting of a deletion eliminating the first two exons of the gene and causing the desired, white-seeded phenotype in this class.

Markers were successfully screened for the candidate mutations in *PvTT8, PvTTG1*, and *PvMYC1* across Black Nightfall, Orca, and a panel of 49 genetically distinct BXO lines. For all three genes, complete co-segregation was found between the genotype at the candidate mutations and color pattern phenotype. This co-segregation was found in all 16 recombinant RILs for *P*, in 9 for *T*, and in 5 for *Bip*. For *Bip*, marker screening and recombinants also removed from consideration the MYB Phvul.010G096400 gene model and all genes further upstream due to recombination in BXO140, and Phvul.010G100300 and genes further downstream based on the WGS data of BXO075. This limited the Bip interval to 39 genes, including *PvMYC1*.

### Gene expression and *PvTT8* intron splicing

In seeds sampled 20 days after anthesis, the expression of Phvul.007G171333 (*PvTT8, P,* Fig. 6) differed significantly between phenotypic groups (*P*=0.002, ANOVA). *P* expression was significantly higher in types with wild-type *P* and *T* (phenotypic class 1, full-colored black seeds) than in any phenotypic class with the *p^hbw^* partial loss-of-function allele (phenotypic classes 4-6; Fisher’s Least Significant Difference test, α = 0.05). With dominant *T*, the *p^hbw^* intron deletion is associated with a 19.5-fold reduction in *P* expression relative to types without the deletion (Fig. 6). *P* expression in *P T* genotypes was also 8-fold higher and non-overlapping with that of *P t* genotypes. Despite this, *P t* types were intermediate in *P* expression and did not significantly differ from any phenotypic categories in *P* expression. No significant differences in the expression of *T* or *Bip* were identified, nor were any differences in gene expression identified in seed samples collected 10 days after anthesis. Amplification of the *PvTT8* region spanning from exon 2 to exon 3 using cDNA templates found uniform excision of intron 3 across *P* and *p^hbw^* types, indicating that the 612 bp deletion in the second intron observed in Black Nightfall and two other lines does not affect intron splicing.

**Fig. 6.**
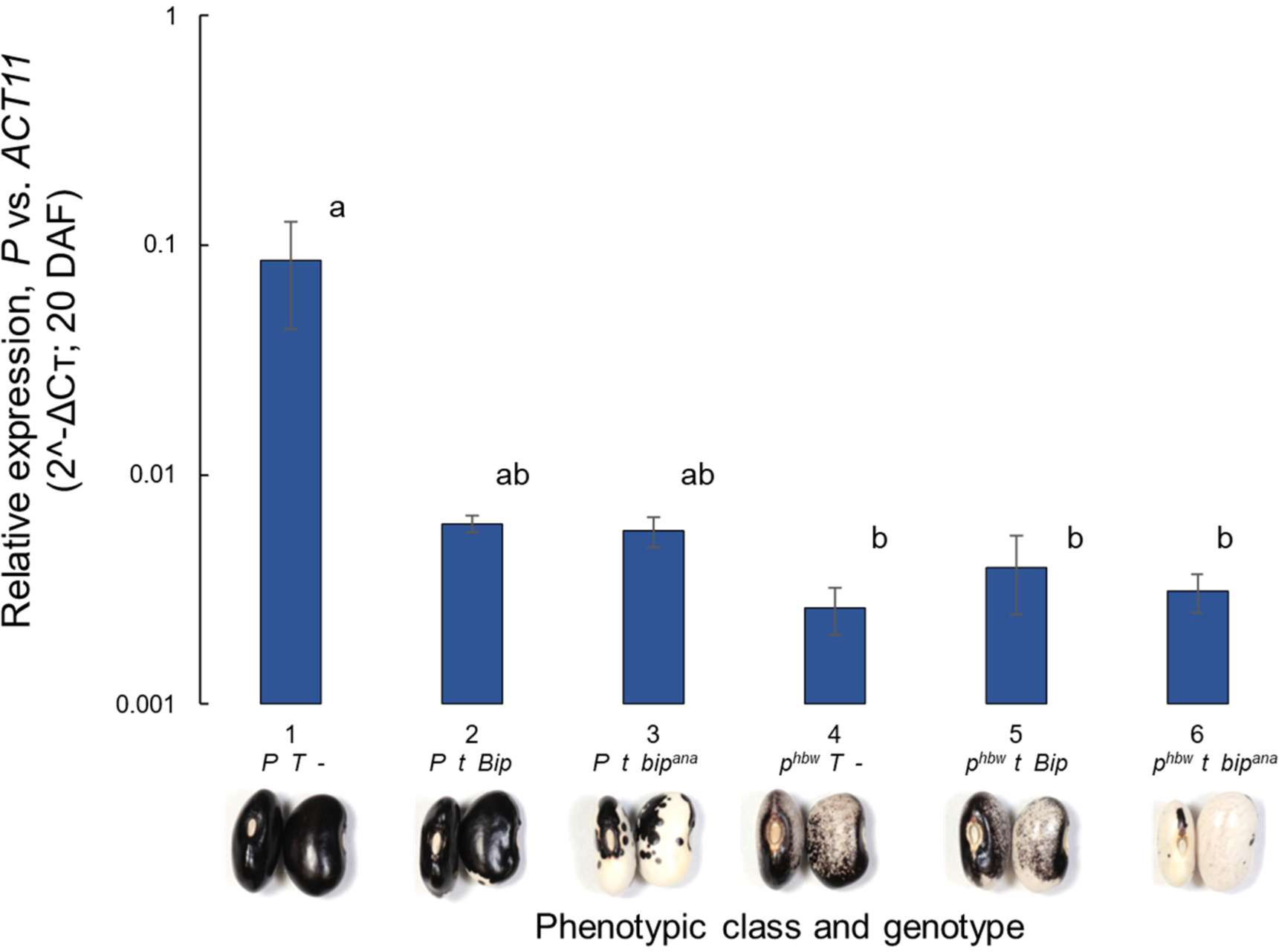
Expression of P differed significantly across BXO phenotypic classes at 20 days after anthesis (P=0.002, ANOVA). RILs with T phbw carry a 612 bp intron deletion in PvTT8 (phenotype class 4) and have 19.5-fold lower P expression than those with P T (phenotype class 1, see Fig. 1). P expression in genotypes with P T (phenotype class 1) was 8-fold higher and nonoverlapping with that of types with P t (phenotype classes 2 and 3), indicating that T may also impact P expression. Bars represent standard error based on three replicates. Significance groups based on Fisher’s Least Significant Difference test (α = 0.05).

### Exploration of candidate genes for *Z* loci

The whole-genome sequencing of 37 diverse accessions allowed for the evaluation of other candidate genes that may control the broad seed color diversity of common bean. BLASTp indicated that common bean gene most similar to the critical Arabidopsis MYB gene *AtTT2* was Phvul.003G132100, *AtTT2* was also the most similar Arabidopsis protein to Phvul.003G132100 (Supporting Fig. S4). Phvul.003G132100 was therefore named *PvTT2.* Phvul.003G132100 is found 1.4 Mb from OAM10^560^, a marker previously mapped 1.4 cM from *Z* (McClean et al. 2002; 31.5 Mb on Pv03). Phvul.003G132100 expression is 22-fold higher in developing seeds compared to any other tissue (O’Rourke et al. 2014; Supporting Figure S5), consistent with a role in the regulation of seed-specific pigmentation. As a candidate gene for *Z*, the whole-genome sequencing data were inspected at this gene model to identify polymorphisms that might be related to seed color patterning.

In 22 Andean accessions, a 315G>T substitution was identified in the Phvul.003G132100 CDS, leading to the substitution of a Lys with an Asn (K105N). This residue is part of the MYB domain of the protein, a region that extends from positions 62-116. The reference genome G19833 (fully-colored Andean genotype) included a Lys at this position, as did the fully-colored Andean variety UCD Canario 707 and all sequenced Middle American genotypes. In contrast, all but one partly-colored Andean types had the Asn at this position. The one partly-colored Andean type without this allele, Vermont Appaloosa, shows the *expansa* pattern consistent with *t Z* (Bassett 2007). To determine the functional importance of the substitution based on sequence conservation, BLASTp was used to align the G19833 amino acid sequence against the NCBI RefSeq protein database. The 100 most similar proteins, spanning 16 plant families, all maintained a Lys at the position. This extremely strong conservation suggests that substitutions at the site have major deleterious effects on protein functionality. These results were further tested using SIFT (https://sift.bii.a-star.edu.sg/www/SIFT_seq_submit2.html). The test results indicated that only Lys would be tolerated at the position (threshold = 0.05, SeqRep = 0.87).

While the variation between *Z* and *z* was mapped to Pv03, the variation between *z* and *z^sel^* has historically been mapped to 1.03 ± 0.33 cM of the *fin* determinacy locus (Bassett 1997). The *fin* locus has subsequently been characterized as *PvTFL1y* (Phvul.001G189200) on Pv01 (Repinski et al. 2012, Kwak et al. 2012), calling into question the localization of genetic factor(s) comprising the reported allelic series for *Z*. Phvul.001G192200, a WD40 repeat protein and close relative of *T*, was identified 245 kb from *fin* on Pv01. Expression patterns of the gene closely resembled those of the *Z* candidate Phvul.003G132100 and *P* (Phvul.007G171333), with expression in developing seeds 41-fold higher than in any non-seed tissue (Supporting Fig. S5, O’Rourke et al. 2014). The four varieties with the *sellatus* pattern (Steuben Yellow Eye, Red Calypso, Snowcap, and Hidatsa Shield; Fig. 4 C6, E3, E4, E6) all showed a single haplotype at Phvul.001G192200, including a deletion of 477 bp from -149 to -641 bp upstream of the TSS. This variant was not found in any other sequenced lines.

### Environmental component of seed pigment distribution

The proportion of pigmented seed coat (PPSC) varied significantly based on both genotype and environment. In the field, the difference in PPSC between varieties was highly significant (*P* < 2.2e-16; Fig. 7), as were the differences based on environment (*P* < 2.2e-16, two-way ANOVA). The interaction between genotype and environment was also highly significant (*P* =2.9e-14, two-way ANOVA). The greatest range in PPSC between environments was seen in the cultivar Orca, which produced pigment on 7±0.3% of the seed coat in the cool climate of Pescadero, CA, compared to 79±5% in the warmer climate of Davis, CA (Fig. 7b). The three varieties with compact growth habit (Orca, UC Southwest Gold, and UC Southwest Red) each had a significantly lower PPSC than any of the four vine types (Zuni Gold, Anasazi, UC Sunrise, UC Four Corners Red; Fisher’s LSD). Location rankings by temperature were the same as by PPSC (Fig. 7b, Supporting Table S3, Supporting Fig. S6). This raised the hypothesis that temperature may have a direct effect on the regulation of pigmentation.

**Fig. 7.**
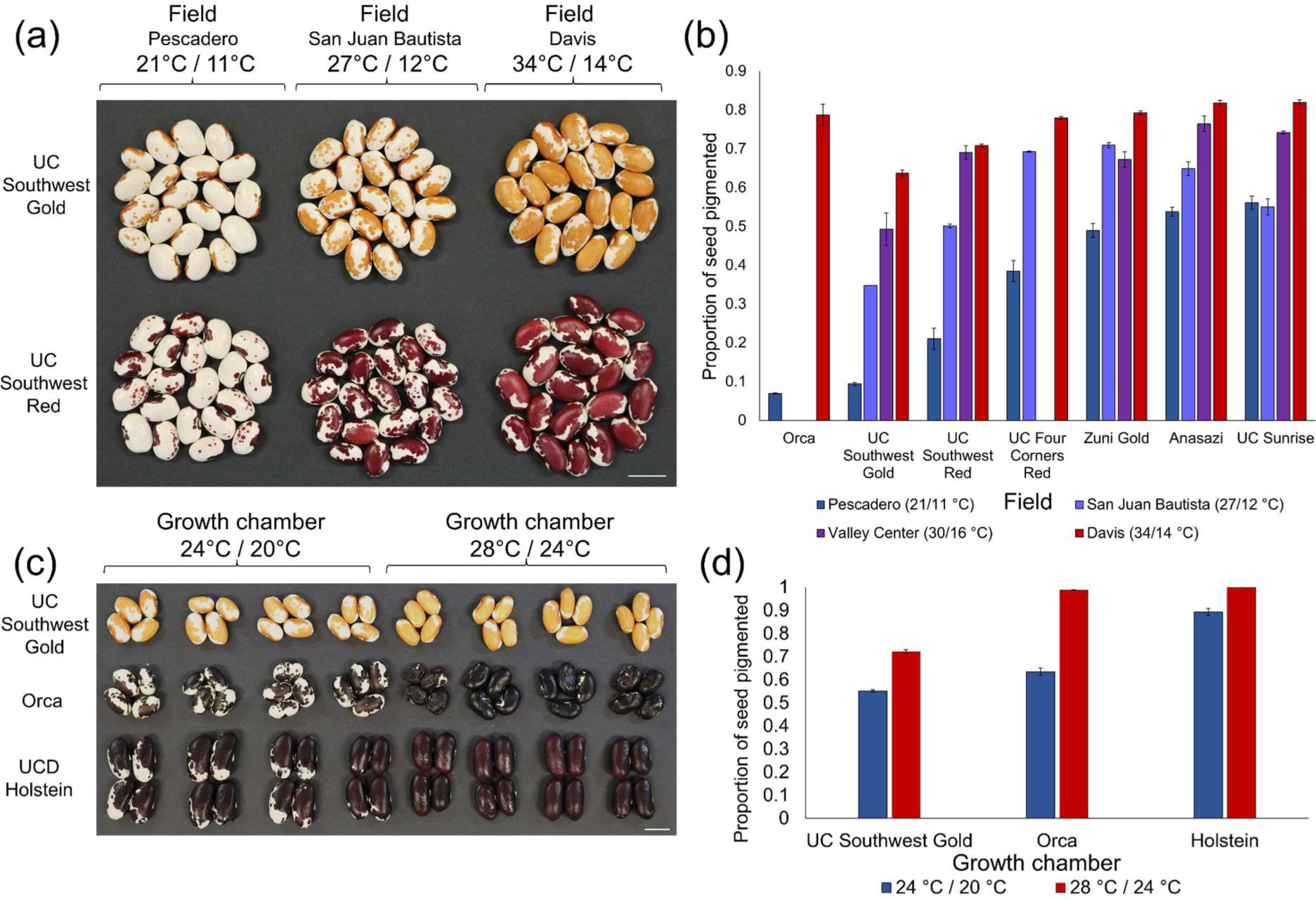
Proportion of pigmented seed coat changes with temperature in field and growth chamber conditions. a) and b) Seeds grown in multi-location field trials (all California, USA) showed significant differences in seed coat pigmentation across environments, with up to 11-fold differences in the proportion of pigmented seed coat between the coolest and warmest environments (July average high/low temperatures shown, °C). c) and d) Growth chamber studies confirm that variation in temperature alone is sufficient to change proportion of seed coat pigmented. A difference of 4 °C was sufficient to cause pigmentation over 20% of the seed coat. Field experiments: 3 bulk-harvested plots per genotype per location; growth chamber: 4 individually harvested plants per genotype per temperature. Error bars represent standard error of the mean. Scale bars are 1 cm.

In identical growth chamber conditions varying only in temperature, the effect of temperature on the PPSC was highly significant (*P* = 9.33e-08, two-way ANOVA), as was the effect of genotype (P = 2.65e-09, two-way ANOVA) and genotype by environment interaction (*P* = 0.002, two-way ANOVA). A temperature difference of just 4 °C between the chambers was sufficient to cause pigmentation over an additional 20% of seed area on average across varieties (Fig. 7d). Seeds (P = 0.002, two-way ANOVA). A temperature difference of just 4 °C between the chambers was sufficient to cause pigmentation over an additional 20% of seed area on average across varieties (Fig. 7d). Seeds grown in the cooler chamber produced pigmentation on 71% of the seed coat, whereas 91% of the seed area was pigmented in the warmer chamber (Supporting Table S4).

## Discussion

### Genetic effects

Our mapping and sequencing results strongly indicate that a series of mutations have occurred within each component of the MBW complexes and that these are responsible for a broad array of color patterns in common bean. In *PvTTG1*, a series of at least seven parallel mutations underpin the loss of function at the *T* locus and are found in all types with partially white seed coats in the species. The identification of these seven separate mutations among the 22 partly-colored accessions in our panel is compelling evidence for strong evolutionary parallelism for the control of seed pattern diversification. These findings coincide with parallel molecular evolution described for other loci in the species, such as *PvTFL1y* (Kwak et al. 2012), *PvTT8* (McClean et al. 2018), and *PvPHYA* (Weller et al. 2019). This evolutionary parallelism also extends to cowpea, in which mutations in the *TTG1* homolog (Vigun09g139900) and secondary genes are required for partial white seed color (Herniter et al. 2019). This is an example of the molecular basis of the law of homologous variation between species (Vavilov 1922).

Our results also indicate that *PvMYC1* likely underlies *Bip*. The *bip^ana^* C217F substitution identified in *PvMYC1* was found in all sequenced partly-colored Middle American varieties, whereas no mutations were identified in the gene in Andean lines. These instead demonstrated mutations at *PvTT2*. The C217F *PvMYC1* substitution occurs in a domain that interacts with WD40 proteins in the related protein *AtTT8* (Zhai et al. 2019). Since the effects of *bip^ana^* are visible only with recessive *t* (a WD40 protein), the *bip^ana^* mutation could achieve its effect due to poor complex formation with the *t^Orca^* protein variant, with poor binding affinity exacerbated by low temperature. Alternatively, in the absence of a functional *T*, *bip^ana^* may be poorer at binding with alternate WD40 proteins to rescue pigmentation. These alternatives could be evaluated through hybridization with varieties like Dapple Grey, which lacks *PvTTG1* in its entirety. The existence of varieties with a full loss-of-function allele *bip* (Bassett 2007) and their difference in phenotype compared to lines with the *bip^ana^* allele indicates that the *bip^ana^*allele likely retains some functionality.

Our characterization of *p^hbw^* sheds light on the partial loss-of-function allelic series at the *P* locus. The conservation of specific motifs in intron 2, as well as the repeated loss of these motifs in partly-colored accessions across three species, and expression differences between *P* alleles, together strongly indicate that this intron contains functionally significant regulatory sequences. The identification of overlapping intron deletions in the homologous genes of partly-colored lima bean (Pl07G000031200) and cowpea (Vigun07g110700) indicates that these may be important for color pattern regulation in these related species as well.

Our sequencing results also shed light on the nature of *Z*. Here, we propose a two-gene model for the *Z*-associated phenotype instead of an allelic series. The identification of two further candidate gene models closely related to *TTG1* (Phvul.001G192200) and *TT2* (Phvul.003G132100), each located in close proximity to mapped “*Z*” alleles, each with nearly seed-specific expression, and each with polymorphisms relating to phenotypic patterns, together makes these compelling candidates for the control of *Z-*related seed color expression. According to a single-gene model, types such as Steuben Yellow Eye (*sellatus* pattern) and Improved Kidney Wax (*virgarcus* pattern) would be expected to differ at *Z* (Bassett 2007, Ernest et al. 2005). Our sequencing results, however, found a single identical mutation in both lines. These *sellatus* and *virgarcus* types were, however, differentiated by the 477 bp deletion upstream of Phvul.001G192200, which is tightly linked to the *PvTFL1y* locus tied to this variation previously (Bassett 1997). We propose to call this Pv01 locus, hypostatic to *Z*, *Sellatus*. A two-gene model for *Z*, rather than an allelic series at a single locus, would parallel the recent breakdown of allelic series at both *bc-1* and *bc-2*, also in common bean (Soler-Garzón et al. 2021a,b).

### Expression and intron splicing

Our expression analyses indicate that the loss of the 612 bp intron fragment in *p^hbw^* types corresponds with a 19.5-fold reduction in *P* transcription. The critical role of intron motifs in promoting transcription is becoming increasingly clear; for many genes, intron motifs are more important for transcriptional regulation than sequences upstream of the TSS (Rose 2019). The non-overlapping and 8-fold lower expression of *P* in genotypes with *P t^orca^* indicates that *T* may also regulate *P* expression, as proposed before the genomics era by Bassett (2003). This fits with the pattern of Arabidopsis, in which *TT8* expression—but not expression of *TTG1* or *MYC1*—is known to be self-reinforced through a positive feedback mechanism by MBW complexes (Baudry et al. 2006). The lack of differential expression of *T* and *Bip* also fits with our sequencing results, in which the *t* and *bip^ana^* mutations of the BXO population affect protein sequence rather than protein abundance. The finding of significant expression differences for *P* only at the later sampling time point, together with the seed coloration patterns of single, double, and triple mutants, is consistent with a model in which the MBW complexes of *Phaseolus vulgaris* propagate pigmentation from the chalaza through the rest of the seed coat. This parallels what has been identified in the development of Arabidopsis seeds (Baudry et al. 2006) and phenotypic descriptions of patterns observed in cowpea (Herniter et al. 2019).

### Environmental effects

Our field and growth chamber studies show compelling evidence that temperature affects the MBW complexes that regulate pigment biosynthesis in partly-colored varieties. In field conditions, the PPSC in Orca varied by 70% between the warmest and coolest conditions, suggesting that activation of flavonoid biosynthesis might be disrupted at low temperatures. These results were confirmed in carefully controlled growth chamber conditions, which isolated temperature as the sole variable. In these conditions, a difference of 4 °C was sufficient to cause a 20% difference in the PPSC, in line with the findings of the field-based studies. This instability may be a hindrance to the large-scale commercialization of partly white-seeded varieties, which are characteristic of some heirloom or specialty bean types but which have been poorly incorporated into large-scale commercial operations where high uniformity is expected. The higher stability of vine-type varieties across temperatures may be explained by differences in pod placement, with pods developing closer to the soil surface and at higher temperatures than those of upright bush types.

Temperature has a strong effect on protein-protein interactions (e.g., Xie et al. 2012). The strong environmental effects of *t*-related seed coat mottling are consistent with a model in which low temperatures disrupt MBW complex activity, causing a failure of signal propagation throughout the seed coat (Fig. 8). This is also consistent with the localization of the putative mutations in *PvMYC1*’s WD40-interacting domain, as well as deletion of part of a WD40 repeat domain in Orca. Alternatively, temperature may affect transcription directly through differential binding of transcription factors to regulatory sequence or other means. Environmental testing was conducted with types carrying *t^orca^ bip^ana^*or *t^cf^ z*, but similar evaluations could be conducted with other mutations at *t, bip*, and *z* to determine whether other alleles (and combinations thereof) would display the same patterns of environmental instability.

**Fig. 8.**
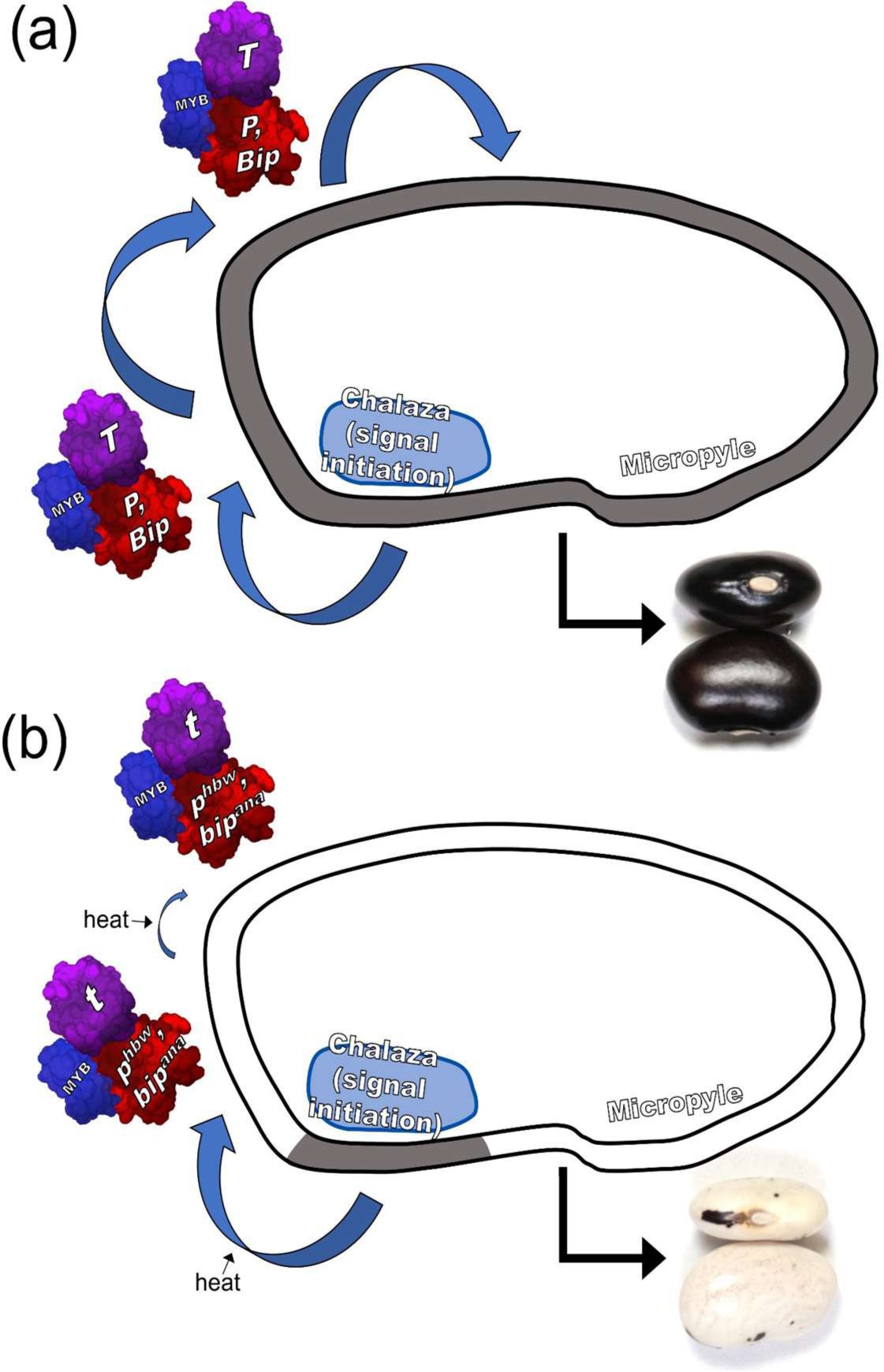
Model for MBW complex-mediated seed color propagation in *Phaseolus vulgaris*. Signal initiation is known to initiate in the chalaza in Arabidopsis (Baudry et al 2006, Xu et al 2015), and propagate from there by MBW complexes through a self-reinforcing signaling mechanism. In a) wild-type plants of common bean (*T*, *P*, *Bip*), this process leads to phenolic pigment production throughout the seed coat. In b) mutant common beans (*t*, *p^hbw^*, and/or *bip^ana^*), this signal propagation does not continue effectively beyond the chalaza (on the left of the hilum in the seeds shown), although higher temperatures appear to correspond to greater propagation. (In all the seeds shown, the micropyle is located on the right of the hilum).

### Parallelism and conclusions

In the BXO population, triple mutants produce pigment only at the chalazal end of the seed, where the patterning signal initiates in Arabidopsis. Lines with MBW mutations fail to propagate these signals at various points across the seed coat (Fig. 1). Our results suggest that proteins forming MBW complexes, including *PvTTG1*, *PvMYC1*, and *PvTT8,* are the fundamental propagators of the pigmentation developmental program throughout the seed coat in common bean.

These results complement those of this study, showing that common bean seed pattern regulation is determined by interactions between each MBW complex component, with or without interactions with temperature. Other related members of these complexes, including Phvul.003G132100 (*PvTT2*) and Phvul.001G192200 (WDR), will be strong candidates for further detailed exploration (Kavas et al. 2022, Supporting Figs. S).

The emerging picture is that genes modifying spatial patterns of pigment deposition—as opposed to pigment composition—appear to encode transcription factors of the MBW complexes. This applies to *T* (*PvTTG1*), *Bip* (*PvMYC1*), *P* (*PvTT8*), *C* (*PvMYB113*), and J (*PvMYB*) and perhaps others such as *Z* (*PvTT2*), and *Sellatus* (*WDR*). Lines carrying mutations at these regulatory genes maintain the metabolic capacity to produce pigments, but the activation of flavonoid pathways does not occur in all cells of the seed coat nor flowers. This stands in contrast to genes controlling pigmentation composition uniformly throughout the seed. These are described as ‘color-modifying genes’ (Bassett 2007) and may correspond to late biosynthetic genes (Xu et al. 2015, Reinprecht et al. 2013). *Violet* (*V*), for example, is encoded by Phvul.006G018800 (flavonoid 3′5′ hydroxylase; García-Fernández et al. 2021, McClean et al. 2022). Other color-modifying genes of this category include *B, G,* and *Rk* (Bassett 2007).

The unique alleles and genes described for the first time in this study are key regulators of seed color and pattern formation, and their encoded proteins are predicted to work together in MBW complexes to achieve their effects. Mutations have occurred at each component of these complexes, sometimes in a highly parallel fashion across different ecogeographic races, gene pools, species, and genera. The interactions between mutations in these complexes gives rise to the broad seed diversity found in common bean. Further, this work has also identified environmental instability across partly-colored varieties of several distinct genetic backgrounds and has shed light on mutations and germplasms that may provide greater color stability. In the future, it will be interesting to compare plant and animal pigmentation and pigmentation pattern and the effect of the environment (Kumar et al. 2022; Elkin et al. 2023; Kratochwil et al. 2023). Deployment of these genetic/genomic resources will be a major step towards maximizing consumer appeal of this important staple crop.

## Acknowledgements

This work was financially supported by grants from the Kirkhouse Trust SCIO and the Organic Farming Research Foundation. The plant material is based upon work supported by the U.S. Department of Agriculture Pulse Crop Health Initiative under Agreement No. 58-3060-1-036. Any opinions, findings, conclusion, or recommendations expressed in this publication are those of the author(s) and do not necessarily reflect the view of the U.S. Department of Agriculture. Talissa Floriani Zimmerman assisted with growth chamber studies. JM, MR, MI, and JA helped manage field trials. Steve Temple developed the unreleased “UCD Jacob’s Cattle” and “UCD Holstein”. Eduardo Blumwald generously provided access to equipment.

## Author contributions

Initial observations and hypothesis generation: TP, PG

Funding acquisition: TP, PG, CP, LT, GB

Project oversight: TP, PG, CD, GB, LT

Experimental methods: TP, TB, TW, VP, MM, AP, CNJ, MRW, SL, SH, AF

Statistical and computational analysis: TP, TB, TW, SL, AF

Manuscript writing: first draft: TP, PG; subsequent drafts: TP, PG, GB, LT, and CD

All authors approved the final version of the manuscript.

## Data availability

Data supporting the findings of this study are available in the Supporting Information of this article. The re-sequence data within 1 Mb of flanking SNPs or previously linked markers will be deposited into the National Center of Biotechnology Information (https://www.ncbi.nlm.nih.gov) with accession no. XXX.

## Supporting information

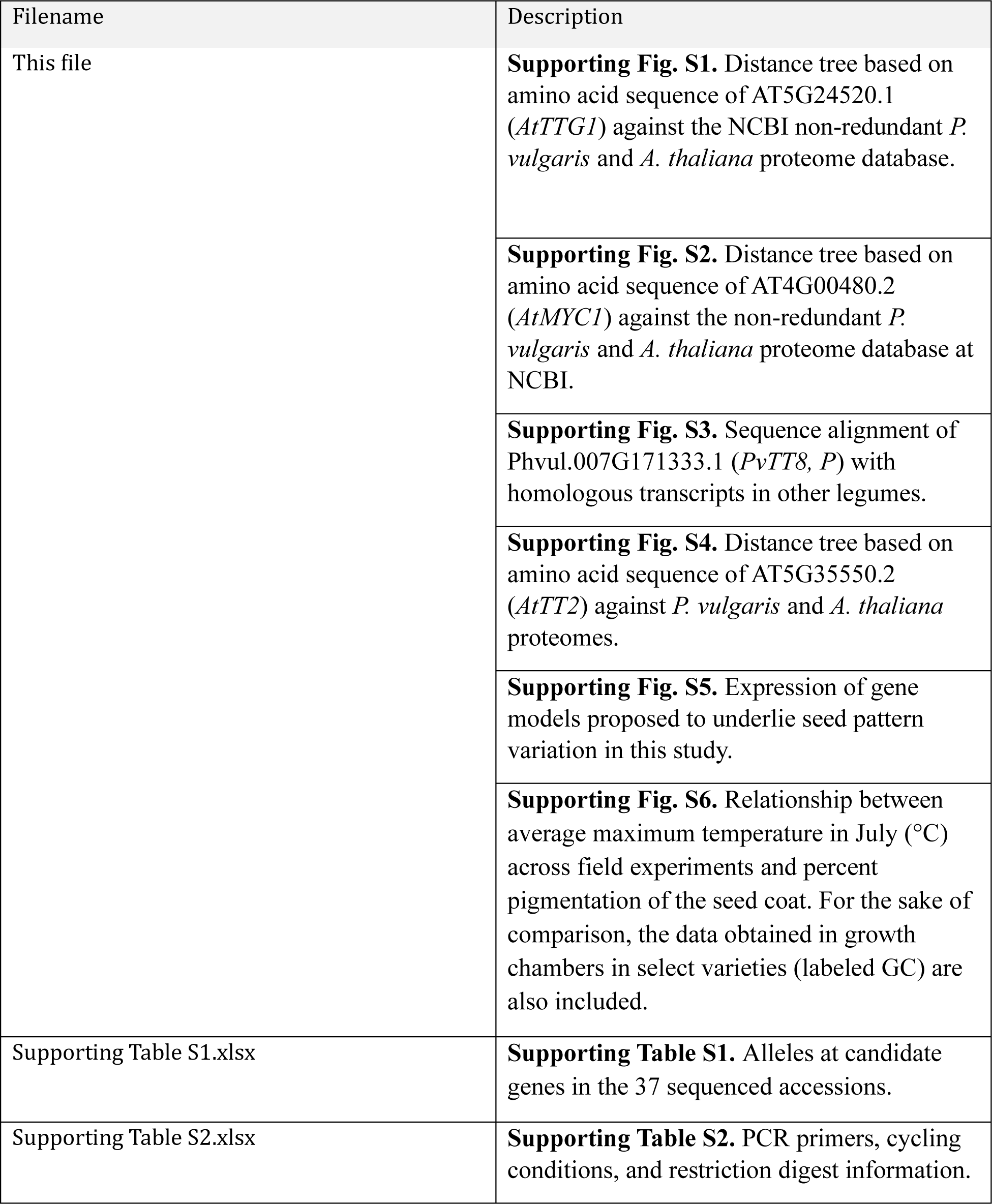
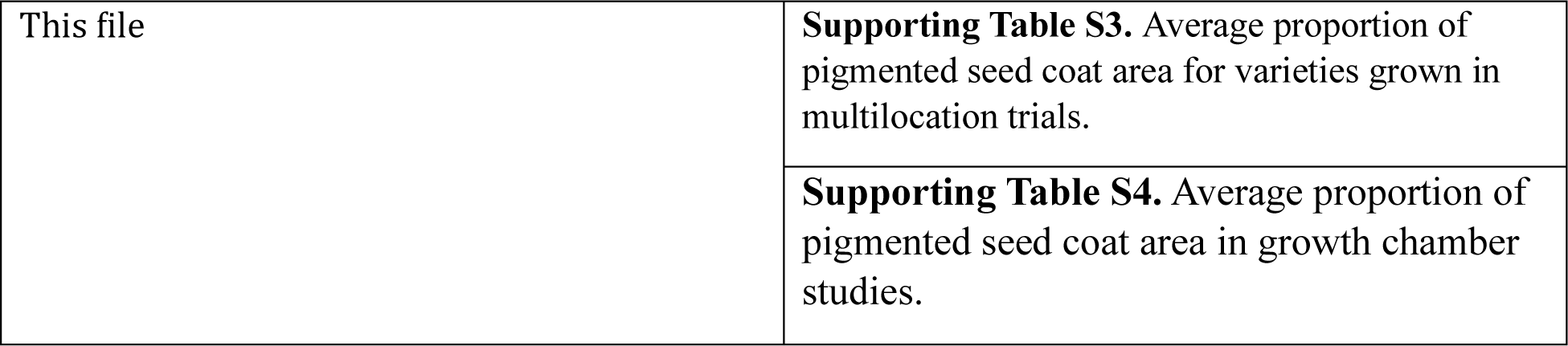

| Filename | Description |
| --- | --- |
| This file | **Supporting Fig. S1.** Distance tree based on amino acid sequence of AT5G24520.1 (*AtTTG1*) against the NCBI non-redundant *P. vulgaris* and *A. thaliana* proteome database. |
|  | **Supporting Fig. S2.** Distance tree based on amino acid sequence of AT4G00480.2 (*AtMYC1*) against the non-redundant *P. vulgaris* and *A. thaliana* proteome database at NCBI. |
|  | **Supporting Fig. S3.** Sequence alignment of Phvul.007G171333.1 (*PvTT8*, *P*) with homologous transcripts in other legumes. |
|  | **Supporting Fig. S4.** Distance tree based on amino acid sequence of AT5G35550.2 (*AtTT2*) against *P. vulgaris* and *A. thaliana* proteomes. |
|  | **Supporting Fig. S5.** Expression of gene models proposed to underlie seed pattern variation in this study. |
|  | **Supporting Fig. S6.** Relationship between average maximum temperature in July (°C) across field experiments and percent pigmentation of the seed coat. For the sake of comparison, the data obtained in growth chambers in select varieties (labeled GC) are also included. |
| Supporting Table S1.xlsx | **Supporting Table S1.** Alleles at candidate genes in the 37 sequenced accessions. |
| Supporting Table S2.xlsx | **Supporting Table S2.** PCR primers, cycling conditions, and restriction digest information. |
| This file | **Supporting Table S3.** Average proportion of pigmented seed coat area for varieties grown in multilocation trials. |
|  | **Supporting Table S4.** Average proportion of pigmented seed coat area in growth chamber studies. |

What is biodiversity?

## Supporting Figures

**Supporting Fig. S1.**
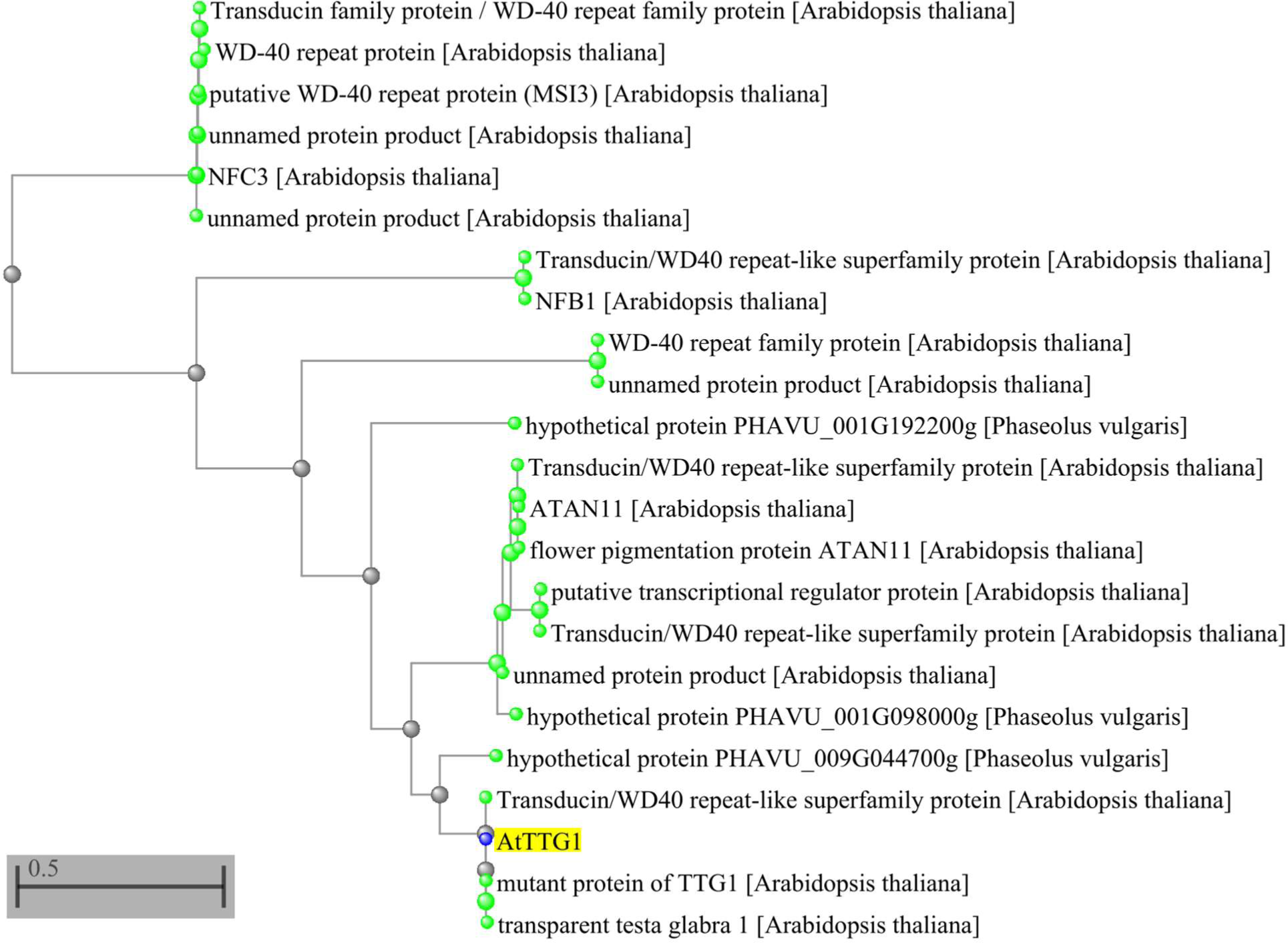
Distance tree based on amino acid sequence of AT5G24520.1 (*AtTTG1*) against the NCBI non-redundant *P. vulgaris* and *A. thaliana* proteome database. Phvul.009G044700 (*T* or *PvTTG1*) clusters most closely with *TTG1* of Arabidopsis, indicating that the two are orthologous sister clades among all proteins of the two species.

**Supporting Fig. S2.**
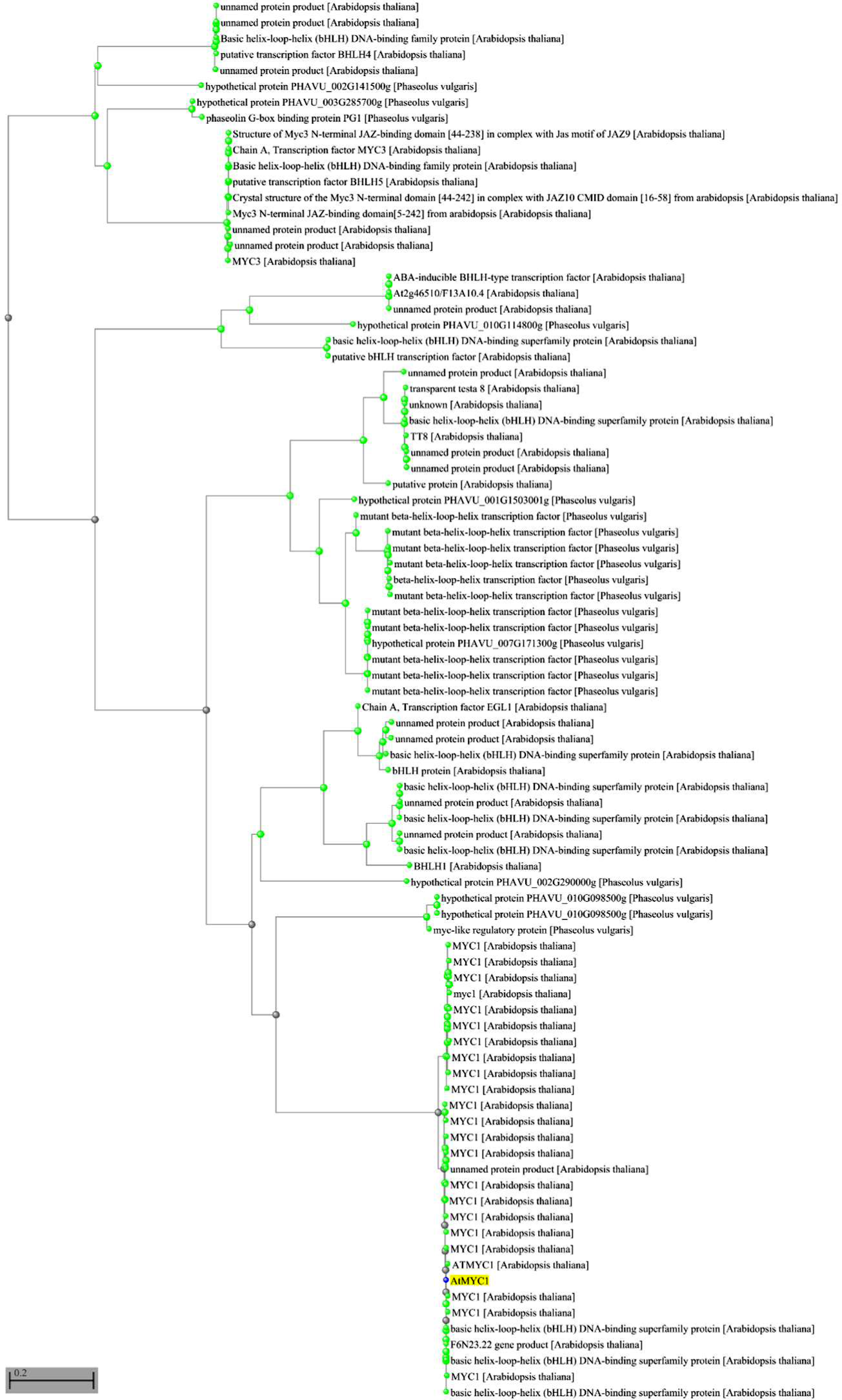
Distance tree based on amino acid sequence of AT4G00480.2 (*AtMYC1*) against the non-redundant *P. vulgaris* and *A. thaliana* proteome database at NCBI. Phvul.010G098500 (*Bip* or *PvMYC1*) clusters most closely with *MYC1* of Arabidopsis, indicating that the two are orthologous sister clades among all proteins of the two species.

**Supporting Fig. S3.**
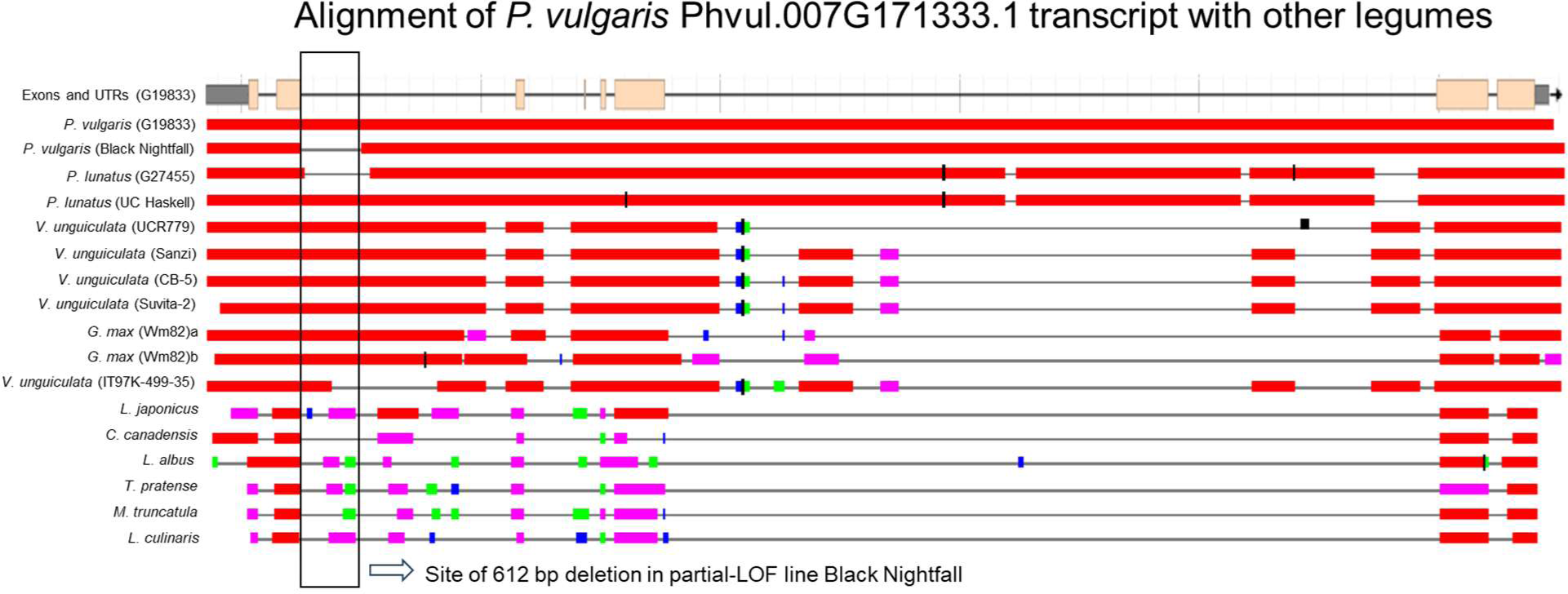
Sequence alignment of Phvul.007G171333.1 (*PvTT8, P*) with homologous transcripts in other legumes. An intron sequence deleted in the partially-colored common bean varieties Black Nightfall and Rio Colorado contains motifs that have been conserved even in distantly related legumes of the cool-season IRLC clade and the basal Papilionoid *Lupinus albus* over at least 40 million years of legume evolution (Koenen et al. 2021). The partly-colored reference lima bean (G27455) and partly-colored reference cowpea (IT97K-499-35) also display deletions in the same intron relative to other varieties of those species with no deletions in the area. The deleted regions of partly colored common bean, lima bean, and cowpea are overlapping.

**Supporting Fig. S4.**
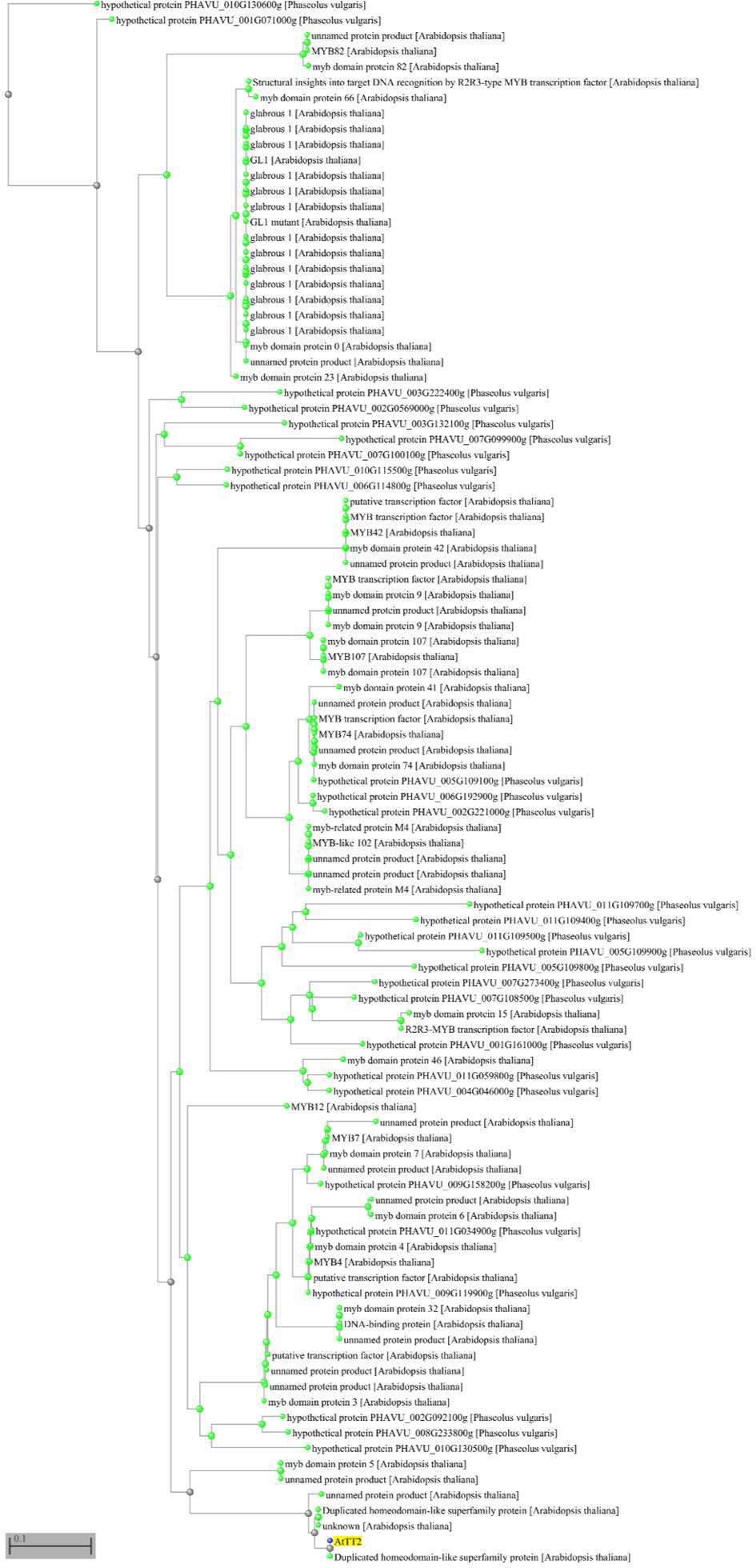
Distance tree based on amino acid sequence of AT5G35550.2 (*AtTT2*) against *P. vulgaris* and *A. thaliana* proteomes. *AtTT2* is a critical MBW complex member and regulator of seed coat pigmentation specification in Arabidopsis. Related genes are strong candidates for control of seed coat pigmentation in common bean. Phvul.010G130600 (at top of figure) is an *AtTT2* relative and was recently shown to underpin the *Joker* (*J*) locus in common bean (Erfatpour and Pauls 2020).

**Supporting Fig. S5.**
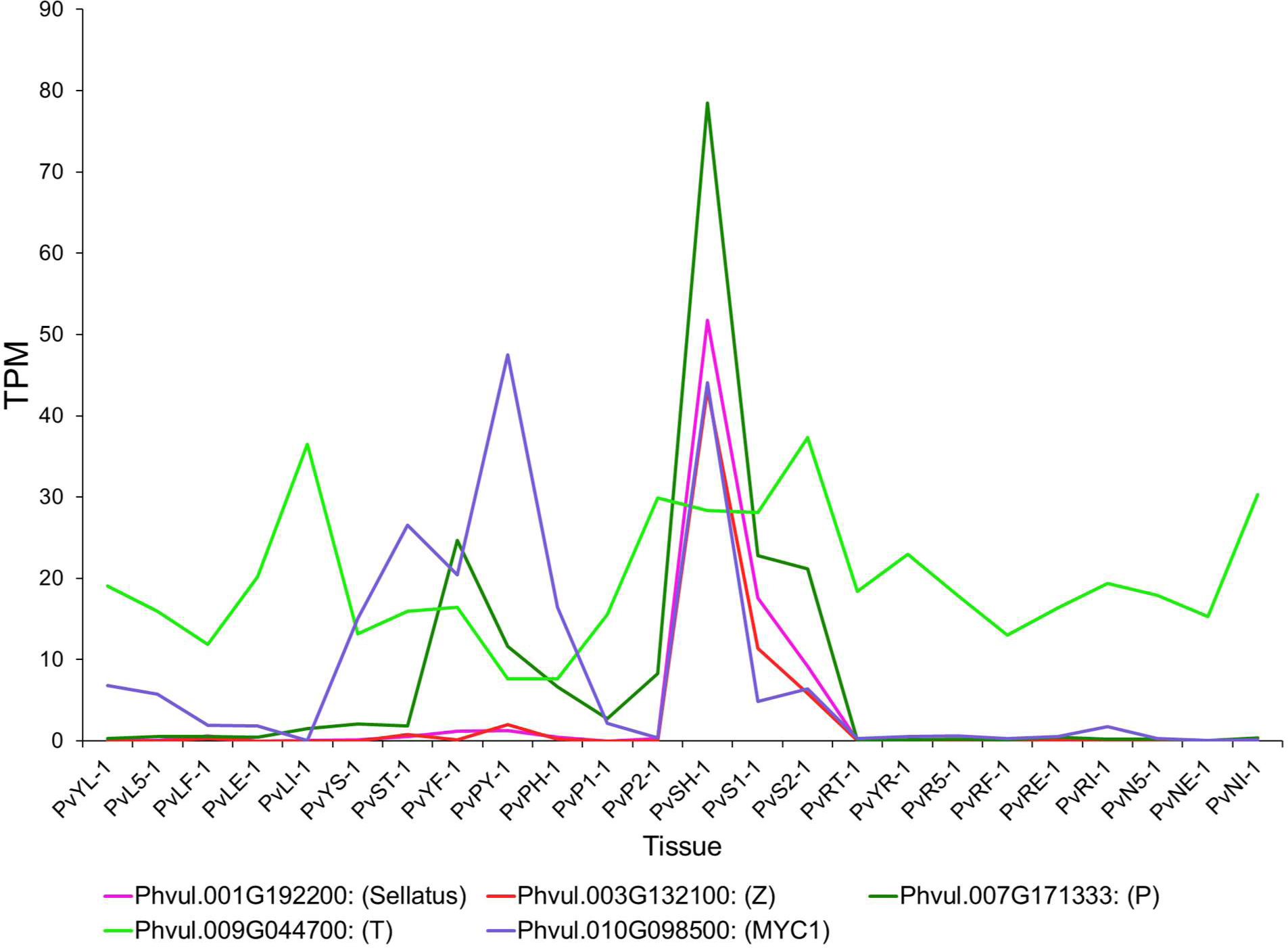
Expression of gene models proposed to underlie seed pattern variation in this study. Data are from O’Rourke et al. (2014) and were collected in the genotype ‘Black Jamapa’ (a Mesoamerican accession). In all cases, very high levels of gene expression occurred in seed development (tissues beginning with ‘PvS’). In some cases, the patterns we-re highly correlated, such as between Phvul.001G192200, Phvul.003G132100, and Phvul.007G171333. PvYL - Fully expanded 2nd trifoliate leaf tissue from plants provided with fertilizer. Tissues: PvL5 - Leaf tissue collected 5 days after plants inoculated with effective rhizobium; PvLF - Leaf tissue, fertilized plants; PvLE - Leaf tissue, plants inoculated with effective rhizobium; PvLI - Leaf tissue, plants inoculated with ineffective rhizobium; PvYS - Young stems; PvST - Shoot tip; PvFY - Young flowers; PvPY - Young pods; PvPH - Pods associated with seeds at heart stage (pod only); PvP1 - Pods associated with stage 1 seeds (pod only); PvP2 - Pods associated with stage 2 seeds (pod only); PvSH - Heart stage seeds; PvS1 - Stage 1 seeds; PvS2 - Stage 2 seeds; PvRT - Root tips; PvYR - Young root material, including tips; PvR5 - Whole roots separated from 5 day old pre-fixing nodules; PvRF - Whole roots from fertilized plants; PvRE - Whole roots separated from fix+ nodules; PvRI - Whole roots separated from fix- nodules; PvN5 - Pre- fixing (effective) nodules collected 5 days after inoculation; PvNE - Effectively fixing nodules; PvNI - Ineffectively fixing nodules.

**Supporting Fig. S6.**
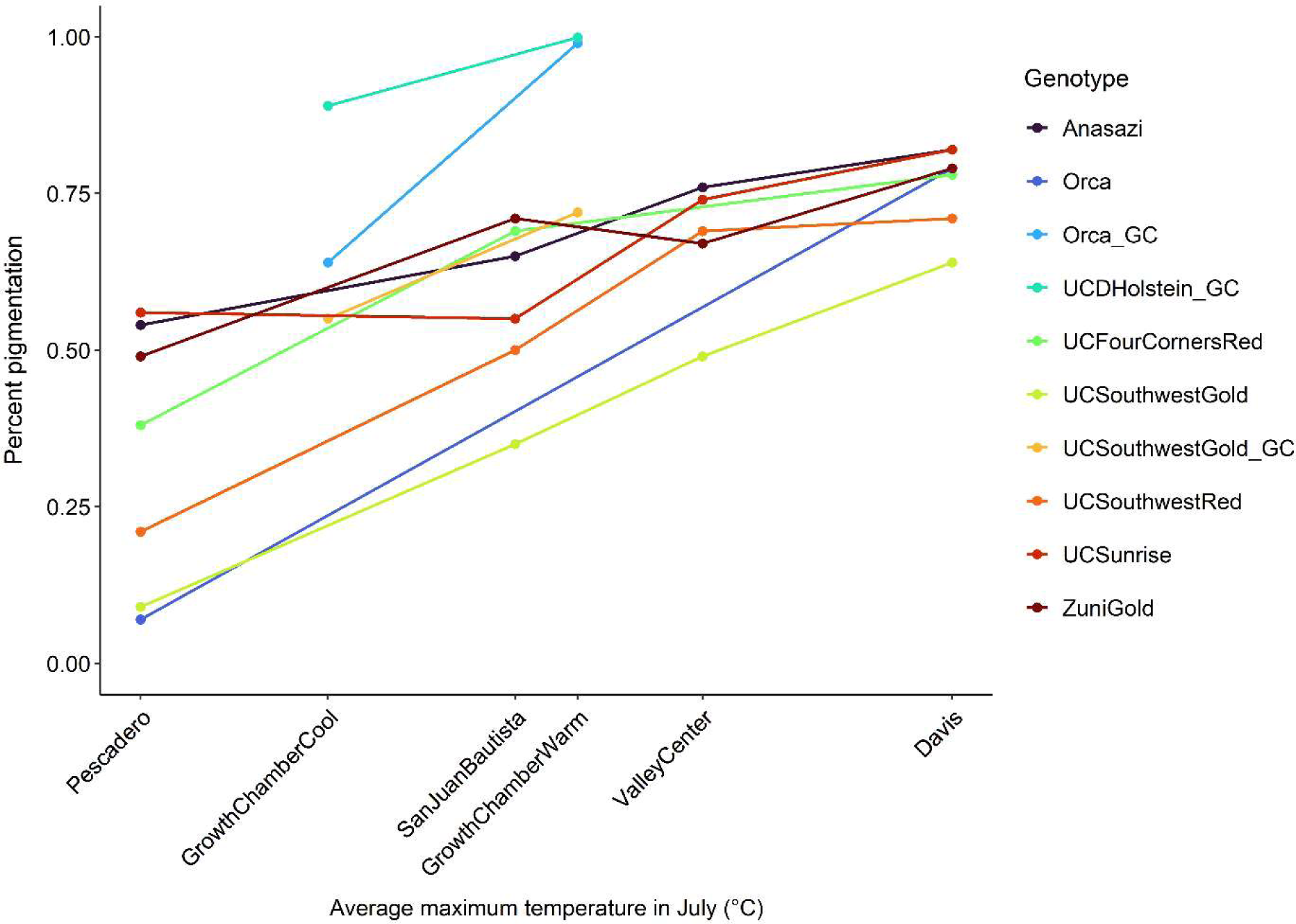
Relationship between average maximum temperature in July (°C) across field experiments and percent pigmentation of the seed coat. For the sake of comparison, the data obtained in growth chambers in select varieties (labeled GC) are also included.

## Supporting Tables

**Supporting Table S1.** Alleles at candidate gene models in the 37 sequenced accessions.

In separate file: Supporting Table S1.xlsx

**Supporting Table S2.** PCR primers, cycling conditions, and restriction digest information.

In separate file: Supporting Table S2.xlsx

**Supporting Table S3.** Average proportion of pigmented seed coat area for varieties grown in multilocation trials.

| Location and<br>July temp. (high/low) | Pescadero | San Juan<br>Bautista | Valley Center | Davis |
| --- | --- | --- | --- | --- |
| Genotype | 21/11 °C | 27/12 °C | 30/16 °C | 34/14 °C |
| Orca | 0.07 | - | - | 0.79 |
| UC Southwest Gold | 0.09 | 0.35 | 0.49 | 0.64 |
| UC Southwest Red | 0.21 | 0.50 | 0.69 | 0.71 |
| UC Four Corners Red | 0.38 | 0.69 | - | 0.78 |
| Zuni Gold | 0.49 | 0.71 | 0.67 | 0.79 |
| Anasazi | 0.54 | 0.65 | 0.76 | 0.82 |
| UC Sunrise | 0.56 | 0.55 | 0.74 | 0.82 |

**Supporting Table S4.** Average proportion of pigmented seed coat area in growth chamber studies.

| Variety | Cool | Warm |
| --- | --- | --- |
|  | 24/20 °C | 28/24 °C |
| UC Southwest Gold | 0.55 | 0.72 |
| Orca | 0.64 | 0.99 |
| UCD Holstein | 0.89 | 0.99 |

